# Softer when Thicker: a Conserved Scaling Law Governs Actin Cortex Mechanics

**DOI:** 10.64898/2026.09.04.749454

**Authors:** Joseph Vermeil, Anumita Jawahar, Valentin Laplaud, Eloise Halouchery, Hugo Lachuer, Camille Plancke, Laura Bernard, Nicolas Borghi, Matthieu Piel, Olivia du Roure, Julien Heuvingh

**Affiliations:** Physique et Mécanique des Milieux Hétérogènes, CNRS, ESPCI Paris, Université PSL, Sorbonne Université, Université Paris Cité, Paris, France; Institut Curie and Institut Pierre Gilles de Gennes, PSL University, CNRS, Paris, France; Institut Jacques Monod, CNRS, Université Paris Cité, Paris, France

## Abstract

Scaling laws are powerful tools in material science to relate geometric and mechanical properties of materials. The actin cortex, a dynamic network beneath the plasma membrane, is essential for cellular morphogenesis, force generation, and tissue organization. Despite its critical role, the mechanical properties of the cortex itself remain poorly understood. Using the magnetic pincher, a new technique we recently developed, we directly measure the thickness *(h)* and elastic modulus *(E)* of the actin cortex in live cells, uncovering a universal scaling *(E ∼ h^−2^)* across multiple cell types. Drawing an analogy to cellular solids, common in nature and material science, we identified an origin for this scaling: the volume fraction of actin filaments varies concomitantly with the cortex thickness, due to a conservation of filamentous actin quantity. Thus thinner cortices exhibit higher actin density and stiffness, while thicker cortices are sparser and softer. The relationship holds under diverse perturbations targeting myosin contractility, actin nucleation, and turnover, but is lost upon actin disassembly, highlighting its dependence on the intact actin network. Our results establish that cells, across cell types, function within a rather limited range of quantity of cortical actin filaments and primarily modulate cortex thickness to tune mechanical properties, with profound implications for cell shape regulation, tissue mechanics, and the interpretation of indentation-based mechanical measurements.

## Introduction

Galileo is credited with the discovery of scaling laws^1^. From the observation that the resistance of a wooden beam scales differently from its length and cross-sectional dimensions, Galileo argued that giants cannot be simple scaled-up versions of humans as they require disproportionately thicker bones, since strength and weight scale differently with size. This insight laid the conceptual foundation for the field of allometry, which examines how biological form and function scale with organism size^2^. Both wood and some types of bones are categorized by material scientists as cellular solids: interconnected networks of solid struts or plates which form the edges or faces of the unit cell^3^. Wood exhibits a hierarchical organization and is structured at the mesoscopic scale into hollow parallel tubes or cells whose wall is made of cellulose fibers embedded in a matrix^4^. Certain bones, like the vertebra and the femoral head have an interior filled with a spongy network of struts, forming an open cellular solid^4^. These cellular solids have unique mechanical properties, one of which being their high mechanical resistance to deformation at low weight^3^. This allows trees and man-made structures such as the Eiffel Tower to reach important heights without buckling on their own weight. This property is usually enclosed in Ashby plots, representing the relation between the modulus E, which quantifies how stiff a material is, and the relative density ρ, the density of the global structure normalized by the density of the struts or wall density, equivalent to the volume fraction. Ashby plots are instrumental for material engineers to select the proper material for an application^5^. On this log-log plot, cellular materials conform to a power-law E∼ρ^α^ where α is generally close to 2. Wood lies in a range of modulus and density that is difficult to reproduce with artificial materials and which has a particularly high efficiency, being able to withstand large stresses with a minimal amount of material.

Some have regarded animal tissues as cellular solids composed of biological cells^6^. The mechanics of the animal cell is controlled by the cytoskeleton, an ensemble of protein fibers arranged in structures of various architectures that are highly heterogeneous in their density. The densest of them is the actin cortex, positioned beneath the plasma membrane and responsible for granting the cell its shape and most of its mechanical properties^7^. The cortex of the cells within a tissue might thus be seen as the equivalent of the walls of a cellular solid^6^. Cell mechanics, however, is largely influenced by the active contractile stress generated by molecular motors in the actin cortex. This contraction generates and hydrostatic pressure resisted by the osmotic pressure through the plasma membrane mechanically coupled to the cortex^8^ and is described as an effective surface tension by analogy with droplet surface tension^9^. This tension manifests itself directly in laser ablation experiments, where the rapid opening of the wound upon cutting illustrates the pre-existing stress in the cortex^10,11^. Spatial gradients in cortical tension are thought to play a fundamental role in cell morphogenesis, driving cell shape changes, polarization and tissue organization^9,12^.

The actin cortex itself is a hollow structure with actin filaments as struts and cellular solids may offer a fruitful framework to understand its mechanics. While traditionally described as a thin, two-dimensional layer of actin filaments, the actin cortex is now recognized as a complex three-dimensional network^13^. Recent advances in super-resolution microscopy, cryo-electron tomography, and expansion microscopy have revealed that the cortex is not a homogeneous layer but a composite network^14^. Actin network architecture results from the dynamic assembly of actin in the presence of numerous partners and regulating factors^15,16^. In the cell cortex, the combination of filament nucleation using formin and Arp2/3, the capping of the growing ends and different disassembly processes regulates filament length distribution^17^. The connectivity is determined by entanglement and cross-linking ensured by proteins such as α-actinin and fascin, as well as molecular motors like myosin II, which generate contractile forces within the network^9,18^. Filament density is mainly set by the balance between nucleation and disassembly, although myosin II-driven compaction of the network may also play a role^19,20^. However, quantifying actin density in the cortex remains experimentally challenging, making it difficult to assess the respective contributions of these processes. In this context, expansion and super-resolution microscopy techniques such as STORM and STED have shed light on the fine structure of the cortex^21–23^, and recent advances in cryo-electron microscopy hold great promise toward a comprehensive, quantitative description of its three-dimensional architecture^24,25^, from which filament density could be directly inferred. The typical mesh size of the actin cortex is considered to be between 20 and 100 nm ^14,26^. This translates into a relative density between 12% and 0.5%, typical of cellular solids. Finally, the actin cortex is a highly dynamic structure, with filaments turning over on a timescale of tens of seconds^9,27^.

Despite its central role in cell mechanics, the mechanics of the cell cortex has so far been studied only indirectly, either using in vitro reconstituted systems or through measurements performed at the whole-cell scale. *In vitro* reconstituted systems have proven crucial to decipher the mechanical properties of actin networks^28–33^. While they have provided a strong basis to frame the possible physical scenarios for cortex mechanics, these studies leave open the question of the actual regimes in which the actin cortex functions in cells. In addition, *in vitro* approaches are still limited in the level of complexity they are able to achieve and often fail to capture the dynamic interactions of the actin cortex with the plasma membrane, linker proteins, and other cytoskeletal components also present in cells. The study of actin cortex mechanics in live cells has long been hindered by the difficulty to access this subcellular structure from the exterior of the cell. Atomic Force Microscopy (AFM) has been routinely used to indent the surface of living cells and measure its mechanical properties^34–39^. These studies either treat the whole cell as an homogenous material to obtain an apparent elastic modulus of the cell, or try to specifically probe the cortex by restricting measurements to shallow indentations with sharp indenters. While the former is useful to compare the overall stiffness of different cells, it gives only indirect information on the cortex, which is much denser than the rest of the cytoskeleton. The latter probes more directly the cortical layer, but the measured elastic modulus still depends critically on unknown parameters, in particular the thickness of the cortex. This is also the case for magnetic twisting cytometry^40^, where a bead attached to the outer layer of the cell is twisted using magnetic torque to probe the mechanics of the cortex, but the interpretation depends on the thickness of the cortex and the bead attachment details. Parallel microplates or AFM with wedged cantilever have been used to deform and flatten whole cells, and these experiments have been instrumental in measuring cortex active tension due to myosin II activity^41–43^. Further deformation allows a measurement of the 2D elastic modulus of the cortex, but its analysis requires idealized assumptions about the shape of the deformed cell^43,44^. As a consequence, a direct measurement of the material properties of the actin cortex layer has proved elusive.

To overcome these limitations, we recently developed the magnetic pincher^45^, which directly measures the geometry and mechanics of the actin cortex in living cells. The technique uses the controlled magnetic attraction between an intracellular and an extracellular superparamagnetic bead to locally compress the cortex while measuring its thickness with nanometric precision. By directly relating the applied force to the resulting deformation, it provides a direct measurement of the cortex elastic modulus without relying on simplifying assumptions. Here, we use the magnetic pincher to demonstrate a robust coupling between actin cortex thickness and elastic modulus at the single cell level and across multiple cell types. Inspired by the mechanics of open cellular solids, we discover that this relationship originates in a E∼ ρ^2^ scaling with conserved amounts of actin. We thus propose that cells, even across cell types, function within a rather limited range of quantity of cortical actin filaments. As a consequence, changes in cortical thickness reflect changes in actin filaments relative density, which directly impacts cortical stiffness. The cell’s control of its cortex mechanics through the control of its thickness could have important consequences on the way cells achieve specific shapes, autonomously and within tissues.

## Results

### Direct probing of the cortex in live cells

To measure the elastic modulus of the cortex in live cells, we modified our magnetic pincher originally developed for thickness measurements^45,46^. This method applies a controlled magnetic force to the cortex of living cells through a pair of magnetic beads, one inside the cell and one outside (Fig. 1 and Supp. Fig. 1). In the initial setup, a low and constant magnetic field held the beads in contact on either side of the cortex to measure its thickness in a time-resolved manner. Here, we dynamically modulated the field to apply compressive forces and derive the mechanical properties.

**Figure 1.**
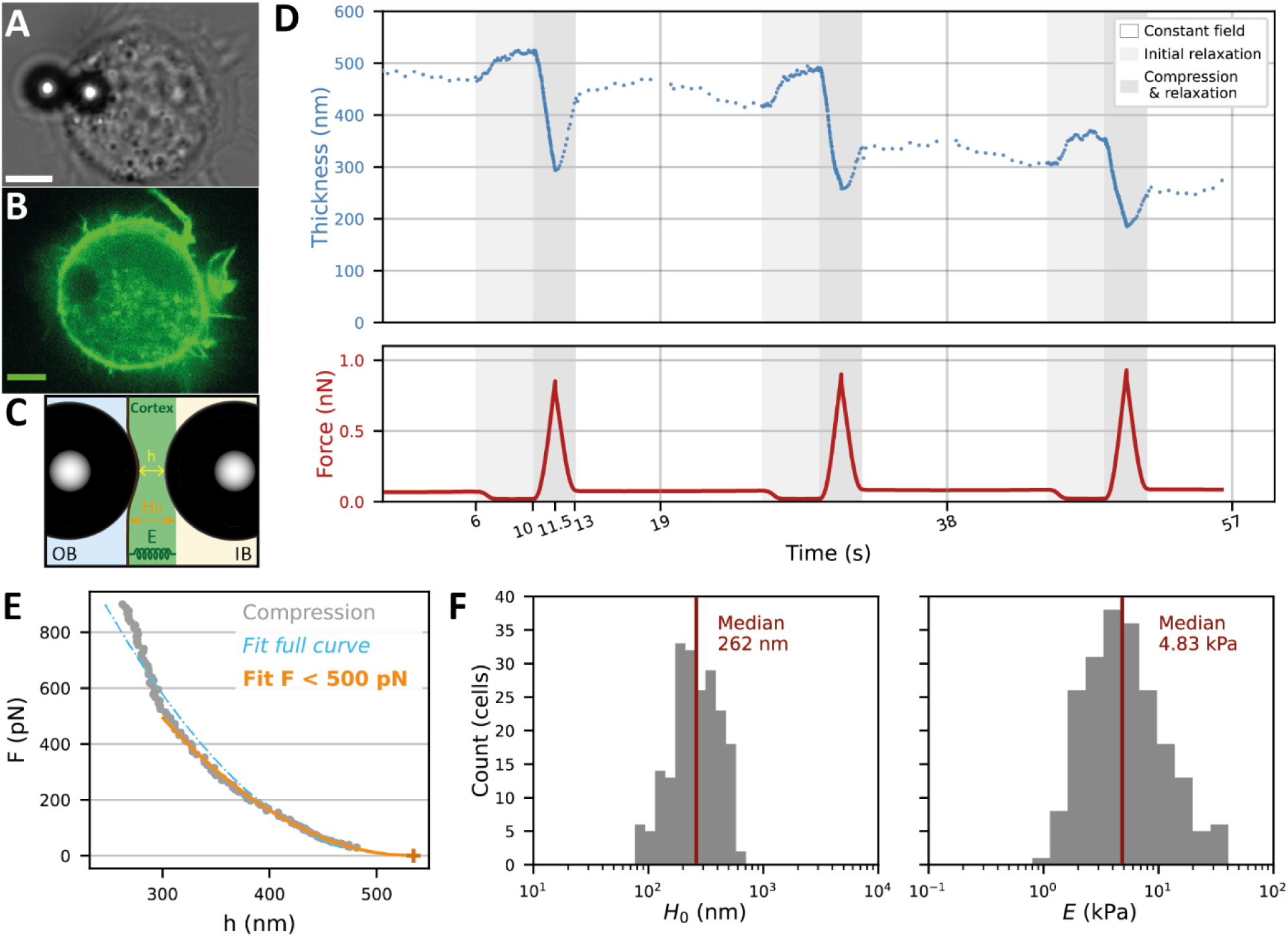
The Magnetic Pincher is a technique to directly probe cortex mechanics in live cells. A, B – Typical configuration to study cortical mechanics with the Magnetic Pincher. 3T3 fibroblast cell adhered on a fibronectin disc has its cortex pinched by a pair of M450 dynabeads oriented in the direction of the external magnetic field (A, Bright field, B, fluorescence image of actin labelled with LifeAct-EGFP, A and B show different cells). C – A schematic view of the measurement at the pinching site. D – Raw data of a typical experiment. Bottom: external magnetic force as a function of time. Top: corresponding cortical thickness. Dark gray areas indicate force peaks (from 1 pN up to about 1 nN), while light gray highlights the short decrease, down to about 20 pN. In between, the resting force corresponds to a constant field of 5 mT. E – Typical example of force-thickness curve (second peak from (D)) and the fitting procedure using the Chadwick model: The fit of the full curve (blue) gives H0 = 516 nm, E = 2.71 kPa. The fit restricted to F < 500 pN (orange) gives H_0_ = 534 nm, E = 2.04 kPa. F – Thickness and stiffness extracted from F<500 pN fits, for the cortex of 3T3 fibroblasts in control conditions (DMSO treated). The x-axes are in log-scale given the log-normal nature of these distributions (see Supp. Fig. 9). Measures were averaged per cell before plotting. N = 1265 compressions on 210 cells from 14 independent experiments.

To introduce beads into the cell, we exploited the phagocytic properties exhibited by most tissue cells, even though they are non-professional phagocytes^47^. Two days prior to the experiments, we incubated 4.5 µm magnetic beads (Dynal M450) with 3T3 fibroblast cells in a culture flask. In this time period, a significant proportion of cells had uptaken one or several beads. We observed increased uptake with fibronectin-coated beads and decreased uptake with PEG-coated beads. We therefore incubated cells with fibronectin-coated beads and used PEG-coated beads externally to prevent uptake during the experiment. Fluorescent microscopy using a live actin dye (LifeAct-EGFP) showed that a shell of actin developed around the bead during uptake but completely disappeared in the course of ∼20s (Supp. Fig. 2).

**Figure 2.**
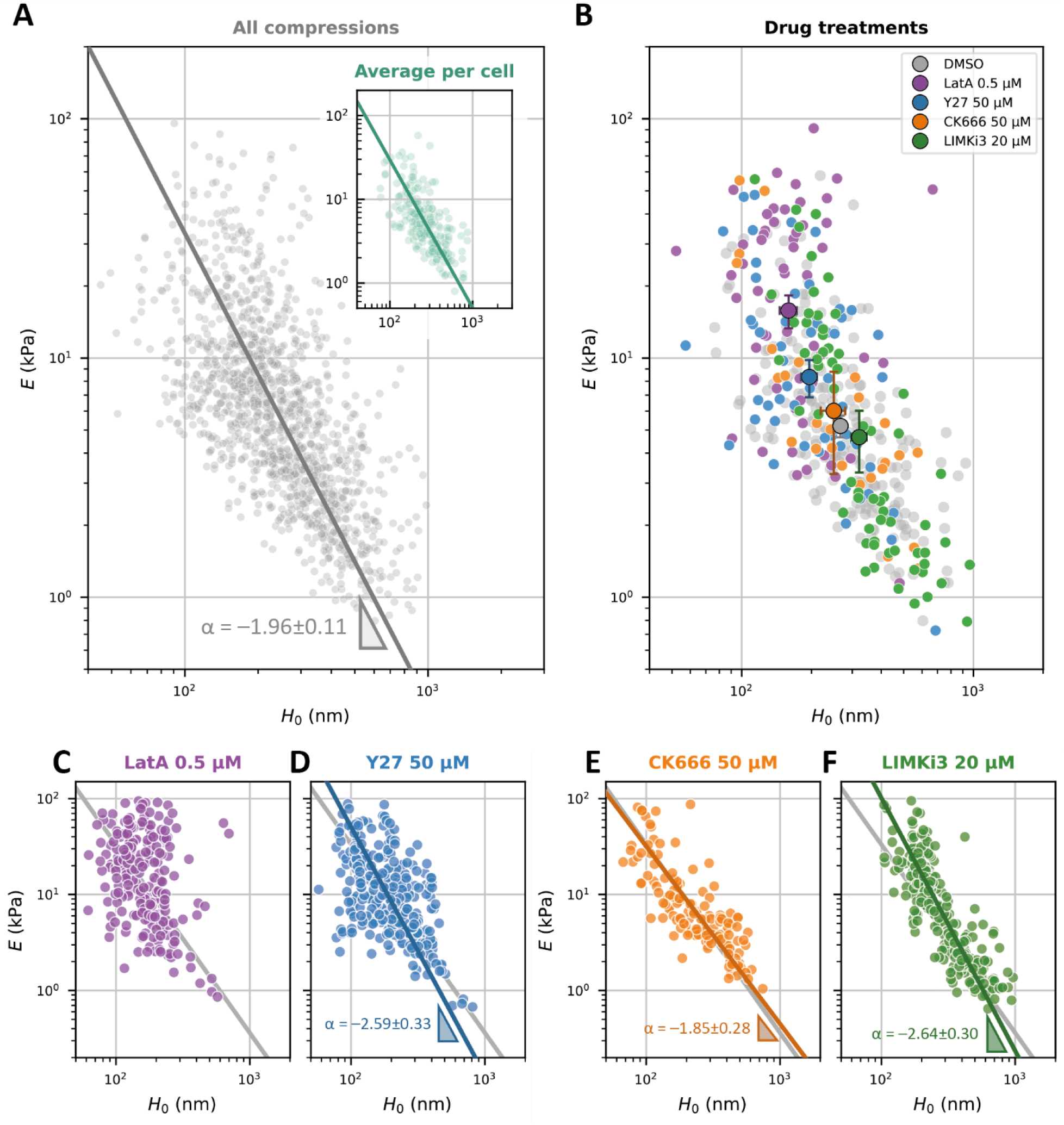
Modulus-thickness scaling of actin cortices. A – Modulus-thickness correlation for 3T3 cortices in control conditions (DMSO treated) for each compression (main graph) and for each cell (inset, average values over a series of compression). Power law fits (gray and green line) were performed using orthogonal distance regression (Pearson’s p-value < 10^-3^). B – Effect of drug treatments on the modulus-thickness scaling. For each drug treatment, the small markers represent the average values per cell, and the large marker with error bars the medians and SEM for H0 and E of treated cells. Light gray points correspond to the control condition, shown as a reference. C-F – For each drug treatment of (B), elastic modulus as a function of thickness for each compression. Corresponding power law fits using orthogonal distance regression (colored line, only if p-value < 0.05). Gray line corresponds to control cells as a reference. Pearson’s p-values and number of replicates for each condition (n compressions, N cells, M experiments): DMSO (control) : p-val < 10^-3^, n = 1265, N = 210, M = 14. LatA : p-val = 0.114, n = 233, N = 57, M = 3. Y27 : p-val < 10^-3^, n = 303, N = 56, M = 3. CK666 : p- val < 10^-3^, n = 137, N = 26, M = 2. LIMKi3 : p-val < 10^-3^, n = 253, N = 65, M = 3.

In order to accurately measure the bead displacement during the cortex deformation, the cell surface at the probing location needs to be roughly perpendicular to the focal plane. Thus, to avoid excessive spreading of the adherent cells, we opted to seed the cells in a chamber patterned with 20 µm-fibronectin discs (Supp. Fig. 3A). Adherent cells formed a near- hemispherical shape, with internalized beads positioned closer to the basal face than to the apical surface (Supp. Fig. 1). Upon introducing PEG-coated beads under a small (5 mT) homogeneous external magnetic field, the dipolar attraction between their magnetic moments led to self-organization into pairs and chains aligned with the field. This set-up generated numerous favorable configurations across the chamber, where the cell surface was sandwiched between bead pairs, either as isolated pairs or as part of chains (see Fig. 1A,B and Supp. Fig. 3B). To prevent internal bead displacement due to cellular movements, we maintained a field of 5 mT corresponding to a holding force of 50–70 pN. The cortex thickness is deduced from the measured distance between the bead centers by subtracting twice the bead radius (see Methods). The precision of the absolute thickness is dominated by the very low polydispersity of the beads (∼30 nm, see Methods), while for cortex deformation, it matches the nanometer-scale accuracy of the distance measurement (see Methods, Supp. Fig. 4 and Laplaud et al.^45^).

**Figure 3.**
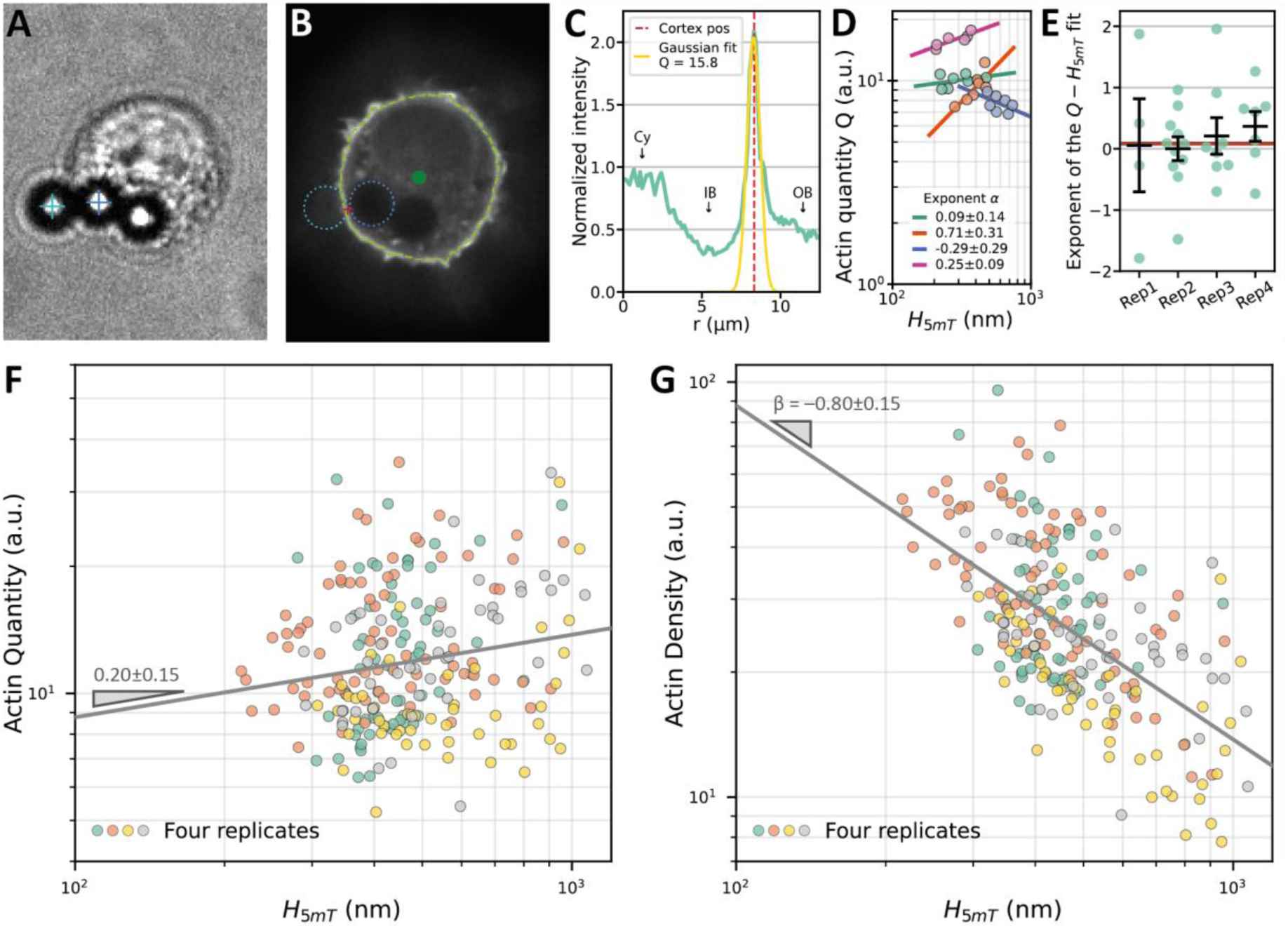
Quantification of actin density in the cortex. A – Cortical thickness was computed from a bright field image using the Magnetic Pincher. B – Confocal fluorescence image obtained using a confocal spinning disk microscope. Image analysis led to detection of the cell center (green dot), the cortex (light green contour), the beads’ position (cyan and blue contours) and the beads’ contact point (red cross). C – Radial profile of normalized intensity from the cell center toward the beads’ contact point. Intensity is normalized by the average intensity in the whole cell. The dashed red line shows the position of the cortex, and the labels “Cy”, “IB” and “OB” indicate the cytoplasm, the inner bead and the outer bead respectively. Cortical actin quantity was computed by fitting a Gaussian on the peak corresponding to the cortex (defined as the Gaussian curve amplitude). D – Typical measurements on 4 cells, from 3 different experiments. Each point corresponds to a paired measurement of the thickness and the actin quantity at the cortex. Each color corresponds to one cell. Exponents were computed by fitting a power law (robust fit with a Huber cost function, 95% confidence intervals). E – Distribution of fitted exponents (as in D) for all cells. The red line shows the median of all points, and the four black points represent the mean and standard deviation from each independent experiment. N = 30 cells from M = 4 experiments. F, G – Actin quantity (F) and density (G) versus thickness (H_5mT_) in log-log scale. Each point corresponds to a paired measurement of the cortical thickness and the actin quantity at the cortex. In (G), density was computed at each time point by dividing the quantity by the thickness. Different colors correspond to different experiments. Exponents were computed by fitting a power law (robust fit with a Huber cost function, 95% confidence intervals). n = 238 points, N = 30 cells, M = 4 experiments.

**Figure 4.**
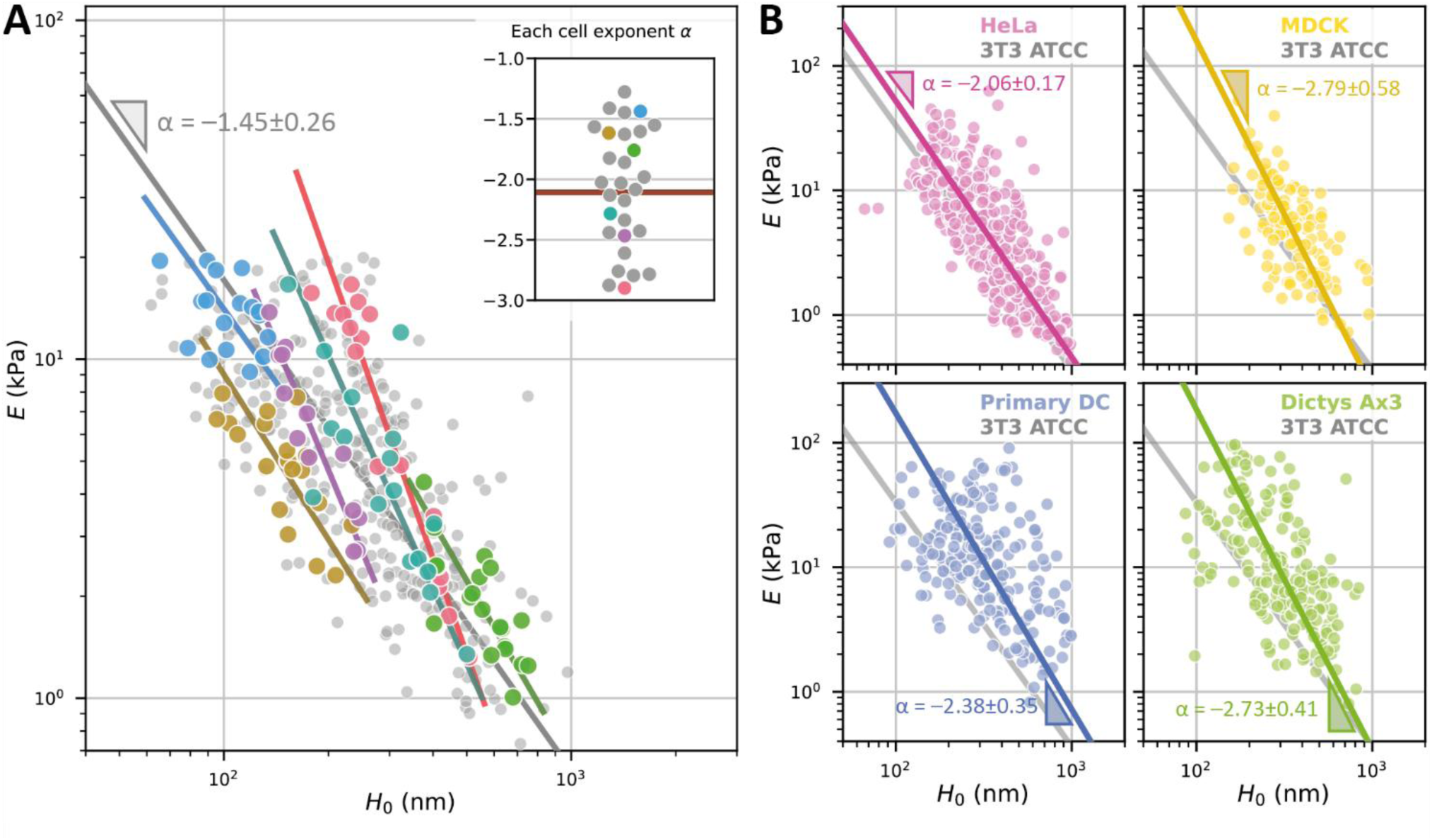
Modulus-thickness coupling exists within single cells temporal evolution and is robust across cell types. A – Modulus-thickness coupling observed in single cells. Cortex indentations are repeated many times over a long period of time (> 3 min) and variations of thickness and stiffness are measured for each cell. Each point represents the thickness and the stiffness from one compression. The 6 colors correspond to 6 different cells. For each cell, the fit of the modulus-thickness coupling is drawn (power-law fit using orthogonal distance regression). The gray line is the power law fit of the whole dataset. In the inset is the distribution of the computed exponents α, with the colored points corresponding to the 6 highlighted cells in the main graph. The median exponent (red line) is -2.11. n = 397 compressions, N = 32 cells, M = 5 experiments. B – Modulus-thickness coupling in diverse cell types. Each point represents the average thickness and the Elastic modulus for one compression. For each cell type the modulus-thickness coupling is fitted with a power-law using orthogonal distance regression. In the background, the light gray line corresponds to 3T3 ATCC cells, shown as a reference. Values for α and 95% confidence intervals are indicated on the fits. For all fits the Pearson’s p-values were lower than 10^-3^. Number of replicates for each condition (n compressions, N cells, M experiments). 3T3 ATCC: n = 1265, N = 210, M = 14. HeLa Fucci cells: n = 463, N = 67, M = 5. MDCK cells: n = 128, N = 26, M = 3. Primary DC: n = 220, N = 22, M = 6. Dictyostelium Ax3: n = 200, N = 29, M = 4.

We applied compressive forces between the beads by modulating the external magnetic field, while simultaneously monitoring the thickness of the indented cortex, in order to generate force-thickness curves and to measure undeformed thickness (H0) and elastic modulus (E) of live cell cortices (Fig. 1C). Low-force data points, essential for extracting those properties from the force-thickness curves, were obtained by temporarily reducing the holding field from 5 mT to 1 mT, thereby lowering the force to 15–20 pN for 4 s to allow for cortex relaxation. We subsequently performed a compression up to ∼1000 pN and a decompression in less than 2 s before reverting to the holding field. Thickness was recorded all along the process with an increased frame rate (100 Hz) during the compression/decompression phase. A typical trace (Fig. 1D) shows an increase of the cortex thickness of ∼50 nm during the initial release of the holding force, followed by an indentation of 200 nm during the compression phase and a recoil to a basal value upon return to the holding force. Five to ten repetitions were performed with a delay of ∼20 s between each. Compression repetitions did not lead to significant change in cortex thickness (Supp. Fig. 5A-B) and it did not show any sign of memory or wearing (Supp. Fig. 5C and 6), consistent with a period between compression phases longer than the characteristic time of actin turnover. Although we observed a consistent hysteresis between an indentation and the following relaxation (Supp. Fig. 7A), we observed no systematic delay between the force maximum and the thickness minimum (Supp. Fig. 7B-C), which indicates that the effective viscosity of the cortex plays a minor role at this timescale.

To determine the cortex’s elastic modulus, we analyzed the force-thickness curves using a Hertzian contact model, routinely used to analyze AFM cell indentation data, but adapted to the magnetic pincher geometry where a thin layer, the cortex, is indented on both sides by two beads whose diameter is much larger than the layer thickness^48^. The situation is assumed to be symmetrical with respect to the cortex median plane and we thus consider half of the geometry. The area of contact between the bead and the cortex is deduced from geometrical argument, allowing for the computation of the mechanical stress (force by unit area). This predicts a contact zone diameter increasing from 300 nm at 20 nm indentation to 1 µm at 200 nm indentation. Because the material’s thickness is equivalent or thinner than the contact zone, the strain should be calculated using the ratio between indentation and thickness (see Methods). In a linear elastic material, stress and strain are proportional through the elastic modulus (E) which can be thus extracted from the force-thickness curve. Most curves are reasonably well fitted by this model, although they sometimes show smaller deformation than predicted at high force (Fig. 1E). This stress-stiffening behavior has been observed in a large number of biopolymer networks^49^, including dense branched actin networks^33,50^. To simplify the analysis, we restricted our analysis to forces below 500 pN, for which stress-stiffening was much less pronounced (see Supp. Fig. 8).

We fitted each force-thickness curve with this model to extract two parameters, the undeformed thickness of the cortex (h0) and its elastic modulus (E) (Fig. 1F). Our method generates large datasets and thus robust statistics (1265 curves for 210 cells). Both thickness and elastic modulus exhibit log-normal distributions (Supp. Fig. 9). Results are thus presented in the following as geometric mean with an interval encompassing one standard deviation. The average thickness of 3T3 fibroblasts is 240 nm (140 - 410 nm), a value similar to what we found previously for dendritic cells^45^. The average elastic modulus is 5.9 kPa (2.4-14.5 kPa), which falls within the range reported in the literature using other techniques (see discussion). In addition, both the undeformed thickness and the elastic modulus were clearly more variable across cells than for successive measurements of the same cell (Supp. Fig. 10), which means that at the timescale of our experiments (3-5 min), the cortex of a cell has a distinct mechanical identity.

### A robust power-law scaling between cortical elastic modulus and thickness

The concomitant measure of the elastic modulus and the thickness allowed us to ask whether cortices of different thickness had different mechanical properties. Plotting the elastic modulus against the thickness, we observed a strong coupling between the two quantities (Fig. 2A): thinner cortices (between 100 and 200 nm) were stiffer (10.2 kPa, (5.0 - 21.0 kPa)), while thick cortices (between 300 nm and 600 nm) were softer (3.1 kPa, (1.6-6.0 kPa)). This inverse correlation was of high certainty (p <10^-4^). A linear regression of these values (n = 1265), revealed a power-law relation between elastic modulus and thickness of E∼h^-1.96±0.22^. The regression bore very similar characteristics when considering the mean of the thickness and elastic modulus for each cell with a relation E∼h^-1.77±0.22^ for cell-averaged values (p < 10^-4^, inset of Fig. 2A).

Next, we verified if the modulus-thickness scaling was robust with alternative metrics. We varied the upper bound in the fit of the force-indentation curves (400 pN, 600 pN, or the full curve). We considered alternative ways to estimate cortical thickness without fitting a model, for instance by measuring its value under the resting field of 5 mT, when the cortex is only slightly indented. Finally we considered an alternative model to fit force-indentation curves, with an expansion of the Hertz contact formula, which could be applied when the indentation is relatively shallow compared to the cortex thickness^51^. With every combination of these metrics for the thickness and the modulus, the correlation was significant with p < 10^-4^ for all combinations and an exponent varying at most of 20%, for all but one metrics (Supp. Fig. 11). This demonstrates the robustness of the modulus-thickness scaling in actin cortices.

### The modulus-thickness scaling is robust to perturbations of the cortex

The cell cortex contains a rich set of proteins regulating its structural properties and its activity. Because we have shown previously that, even when actin filaments are strongly depolymerized, there is still a significant cortical thickness (∼90 nm ^45^), we first assessed the mechanical properties of this remaining layer by treating cells with Latrunculin A (for the dose response effect on cortical actin see Supp. Fig. 12). The treated cells displayed thin and stiff cortices, but the coupling between the two was lost (Pearson’s p-value= 0.114, Fig. 2C), showing that the modulus-thickness scaling is a property of the intact actin cortex. We then perturbed the actin cortex dynamics by targeting an upstream regulators of myosin II motor activity (ROCK, using Y27632) (Fig. 2D). We perturbed the actin cortex structure, using CK666 to inhibit Arp2/3 based branched nucleation (Fig. 2E). We perturbed actin severing and turnover via an upstream regulator of cofilin, LIMK (using LIMKi3, see Fig. 2F), which promotes actin polymerization^52^. All these perturbations changed both the average thickness and elastic modulus of the population of treated cells, moving along the power-law between thickness and elastic modulus of control cells (Fig. 2). For example, although Y27632 treatment led to significantly thinner cortices, control and treated cells with similar thickness had similar elastic modulus (Supp. Fig. 13). Furthermore, contrary to Lat A-treated cells, for all these treatments, the scaling between thickness and elastic modulus was preserved within the population of treated cells, with a power-law relation of E∼h^α^ (α remaining close to -2, see Fig. 2). These experiments demonstrate that the power-law scaling between thickness and elastic modulus, although a property of the actin network, does not depend on the details of its structure or dynamics.

### Cellular solid scaling can explain the coupling between thickness and modulus as a scaling between cortex thickness and actin density

We reasoned that the coupling between the elastic modulus and the thickness of the cortex could result from generic material properties of the actin network. Treating dense actin networks as cellular solids, in which filaments bend between crosslinkers or contact points in the network, predicts a scaling law^3^ where the elastic modulus of the network increases quadratically with the relative density of the actin cortex E∼ρ^2^. The combination of this scaling with our experimental observation that E∼ h^-2^, implies that the cortex thickness should scale as the inverse of the network density (h∼ρ^-1^), which simply reveals that the quantity of filamentous actin in the cortex remains constant between thick and thin cortices. This conservation of the total quantity of actin in the cortex would constitute a simple explanation for the observed coupling and its robustness to perturbations.

To directly test this hypothesis, we completed the measures provided by the magnetic pincher, with a measure of actin fluorescence density near the location of the probing. We used Lifeact-EGFP expressing 3T3 cells and imaged them in 3D using confocal fluorescence microscopy (see Fig. 3A). On an image of cortical actin (see Methods) we selected a location a few hundreds of nm below the contact point, to avoid the shade of the beads. We then computed a radial intensity profile from the cell center, encompassing the cortex at this location and the peak corresponding to the cortex was fitted by a Gaussian curve. The quantity of actin per cortex area was measured as the integration of this curve along the thickness, normalized by the whole cell average to account for the diverse level of expression in LifeAct-EGFP between different cells (see Methods for more details and Supp. Fig. 14). The density of the network (or quantity by volume) was taken as the quantity per area divided by the thickness measured at the same place with the magnetic pincher, about 4 seconds after fluorescence imaging.

The local actin quantity thus measured depended very weakly on the measured thickness of the cortex, neither at the single cell (Fig. 3D-E) nor at the population level (Fig. 3F) and thicker cortices (median above 400 nm) did not contain significantly more actin than thinner ones (see Supp. Fig. 15 A-B). On the contrary, actin density was strongly correlated with the thickness (Fig. 3G), with thinner cortices (smaller than 400 nm) being typically 1.5-times denser than the ticker ones (see Supp. Fig. 15 A-B). The correlation between these quantities is significative (p = 1.8⋅10^-4^), and holds even when taking only the median values for each measured cell (Supp. Fig. 15 C-D). The regression suggests a power-law of ρ∼h^β^ with an exponent β of −0.8 ± 0.15, close to the −1 exponent we anticipated from our simple scaling analysis: a scaling of ρ∼h^-0.8^ with the scaling E ∼ ρ^2^ entails a relation of E∼h^-1.6±0.3^, close to the experimental scaling E∼h^-1.96±0.22^. The small range of actin quantity that we observed experimentally reduces the precision on the power-law exponent in the relation between the density ρ and the thickness h. Nevertheless, our measures strongly suggest that the main parameter that changes between cells displaying cortices of various thicknesses is the density and not the quantity of actin, resulting in thinner cortices being denser and thus stiffer.

### Modulus-thickness scaling in single cells

We previously showed that cortex thickness spontaneously fluctuated in single cells at a timescale of 20 seconds and over a large amplitude, in a myosin-II dependent manner^45^. We thus wondered whether the modulus-thickness scaling held in single cells and performed an experimental series where the cortex of a single cell was measured at least eight times in a course of a few minutes. As expected from the actin conservation data (Fig. 3), a significant correlation (p < 0.05) between elastic modulus and thickness was present in 80 % of the single cells tested (Fig. 4A). The value of the exponent in the power-law relation of E∼h^α^, calculated for each single cell over time, had an average value of -2.31 ± 0.80 and did not significantly differ from the value measured at the population level (-1.96 ± 0.22). The regression lines of individual cells fell either above or below the general regression line for the population measures (Fig. 4A). Assuming invariant structural architecture of the cortices of these individual cells, this would reflect the variation in cortical actin quantity from one cell to the other. This variation can be assessed by calculating the elastic modulus value that these cortices would have at a given thickness using the power-law scaling for each individual cell. We obtained a distribution of an effective elastic modulus for the individual cells, calculated at 300 nm (approximately the mean thickness of the population), with a mean value of 3.0 kPa (1.5 - 6.3 kPa). The geometric standard deviation of this distribution showed a factor of 2.1 in width, suggesting an average variation of 44% in the quantity of cortical filamentous actin between individual 3T3 cells (Supp. Fig. 16). This rather limited variation means that the quantity of cortical actin is under tight control in a population of cells.

### The modulus-thickness scaling is robust across a variety of immortalized and primary cells from different species

Since the conservation of the cortical actin quantity in a population of cultured 3T3 cells appeared tighter than we would have expected, we asked whether this result could also be observed in different cell types, including primary cells and cells from different species. We selected for this goal four other extensively studied cell types: mouse primary immune cells (dendritic cells, DC), the Dictyostelium Discoideum amoeba (dicty), human cervix cancer cells (HeLa cells) and canine kidney epithelial cells (MDCK). All cell types were able to uptake M450 magnetic beads, with an adapted protocol (see Methods and Supp. Fig. 17). Hela and MDCK cells were adhered on 20 µm fibronectin patterns, while the non-adherent DC and dicty cells were simply placed on a BSA-coated glass substrate on which they weakly adhered without spreading. Thickness and stiffness measurements were performed as described above for 3T3 cells.

In all the cell types, a significant correlation was found between cortex thickness and elastic modulus, like in 3T3 cells. The scaling exponent ranged from -2.06 ±0.34 in Hela cells to -2.79 ±1.17 in MDCK and the relation was, in all four cell types, compatible with a power law E∼h^-2^, suggesting that the density variation dominated the scaling between thickness and elastic modulus in these different cell types. Although the median thickness and elastic modulus of the different cell types were significantly different, the variation of the median elastic modulus was mostly explained by the variation of the median thickness: HeLa cells had, on average, a thicker and softer cortex than 3T3 cells, but fell on almost the same modulus-thickness relation (Fig. 4B). This suggests that the quantity of filamentous actin present in the cortices of the two cell types was similar, but that HeLa cells have thicker, softer and less dense cortices than 3T3 cells. In the case of DCs and dicty, the cortex thickness was similar to the one of 3T3 cells (p > 0.05 for both), but significantly stiffer (p<10^-2^, p<10^-3^) (see Supp. Fig. 18). This translates into a regression line that is nearly parallel but shifted upward relative to that of 3T3 cells. This would imply that, despite their distant evolutionary origins, both amoeboid cell types exhibit a similarly elevated quantity of actin filaments at their cortex as compared to 3T3 cells. In conclusion, our analysis of the coupling between thickness and elastic modulus of widely different cell types demonstrates that similar power laws are observed, well explained by a scaling between modulus and density typical of cellular solids and a conservation of the amount of actin filaments in the cortex of a given cell type. In addition, the comparison between cell types reveals that widely different cells, either in terms of tissue or species, might have the same quantity of actin in their cortices when they shared similar phenotypes: in our dataset, amoeboid cells (DC, dicty) appeared to have more cortical actin than mesenchymal cells (3T3, HeLa, MDCK) independently of the species.

## Discussion

### A Universal Scaling Between Cortex Thickness and Elastic modulus

Our study demonstrates a robust and universal power-law relationship between the thickness and elastic modulus of the actin cortex, measured directly in live cells using our magnetic pincher. This relationship, characterized by E∝h^−2^, holds across multiple cell types, including fibroblasts, immune cells, and epithelial cells, and persists even when the cortex is perturbed by drugs targeting myosin contractility or actin regulation factors, but is lost upon actin disassembly, suggesting that it is a property of the cortical actin network. Considering the actin cortex as a cellular solid whose modulus scales with relative density as E∼ρ^2^ can explain this scaling: variations in cortex thickness are accompanied by changes in cortical actin relative density, while the total quantity of actin remains largely conserved. This conclusion is supported by independent measurements of actin fluorescence density, which confirms that thinner cortices are denser and stiffer, while thicker cortices are sparser and softer.

While the magnetic pincher offers a powerful tool to measure cortex mechanics, it requires some choices to determine thickness and elastic modulus from a fitting procedure. We showed, however, that the modulus-thickness scaling unraveled here was not affected by different combinations of alternative metrics, further reinforcing the robustness of this result (Supp. Fig. 11). Our fluorescence-based measurements of actin density clearly support the hypothesis that the coupling between thickness and elastic modulus arises from variations in density. However, these measurements are imperfect as fluorescence was measured close to, but not at the exact location of the thickness measurement. Future studies could employ super-resolution microscopy techniques to visualize actin organization at higher resolution and provide insights into the dynamics of actin network, which could complement our mechanical measurements and offer a more comprehensive understanding of how actin density and organization contribute to cortex mechanics. As previously observed^45^, removing actin through Latrunculin treatment did not result in the total disappearance of the measured boundary layer. This remaining layer may be composed of components such as spectrin or intermediate filaments that have been reported in the cortex^53,54^. The modulus to density scaling observed in this work suggests that these materials would be embedded within the actin cortex rather than forming a separate layer, since the scaling between E and h would differ substantially if two distinct layers were being deformed in series. Identifying the specific components of the boundary layer that remain after actin removal lies beyond the scope of this work.

The cortex may exhibit different mechanical properties parallel and perpendicular to the plasma membrane, an anisotropy that our approach cannot capture. Studying cortical anisotropy will require further technical developments and is out of the scope of this study. Additionally, the magnetic pincher probes only a small region of the cortex, with a diameter ranging from 300 nm for 20 nm indentation to 1 µm for 200 nm indentation. Given the dynamic nature of the actin cortex, with a turnover time of a few tens of seconds, the observed temporal variations of our measurements, over a large number of cells, represent an average behavior that includes the spatial scale, at least in non-polarized cells.

### Modulus density scaling and density thickness scaling

The modulus thickness scaling we discovered here is explained by a combined effect of the modulus density scaling and the density thickness scaling. The modulus density scaling, typical of cellular solids, can be examined in the case of the actin cortex. This relation stems from the bending of actin filaments considered as struts. If we consider the cortex as a network of filaments with a length between contact point L (mesh size), the relation between the stress (σ) and the force (F) acting on one filament is σ=F/L^2^. Because, in the cell cortex, L is much shorter than Lp, the persistence length (∼10µm for actin filaments), we can consider that bending is only enthalpic and stems from network deformation^32^ rather than from entropic fluctuations. The force to bend a filament with an amplitude dL scales as F∼B*dL/L^3^, with B the bending modulus. With the hypothesis that deformation is homogeneous in the network, dL/L=ε the network deformation and F∼B*ε/L^2^. The elastic modulus E, defined by σ= E*ε, thus scales as E∼L^-4^ ^55^. The relative density of the network, ρ, (volume fraction of filaments), and the mesh size, L, depend one on the other, as L∼ ρ^-½^ ^56^ and thus E ∼ ρ^2^.

In vitro, similar scaling has been described for shear modulus in networks of lower relative density. The modulus of entangled and crosslinked actin networks was found to vary with actin concentration consistent with a power law scaling of exponent 1.8 for entangled actin networks^29^, and 2.5 for crosslinked networks^30^. These values are close to the exponent 11/5 predicted from the entropic resistance to elongation of semi-flexible polymers^57^. A modulus density scaling was also reported in vitro dense branched actin networks^33^, with a much shallower exponent of 0.6. In this case, the variation in actin relative density is due to a modification of the growth stress of the network, which could influence the density modulus scaling.

Although a density thickness scaling in the actin cortex has never been formulated as such, the relationship between thickness and actin density of cell cortices has been reported in several studies. Measurements of cortex thickness in live cells is possible with a dual labelling of the cortex and the membrane^58^ and has been occasionally associated with an evaluation of the actin quantity in the cortex^17,59^. This allows for an estimation of quantity and density comparing different situations. In Chugh et al^17^, the cortex of mitotic cells was shown to be thinner and of increased density relative to interphase, with a similar amount of actin, holding true in various cell types: HeLa, HeLa-S, normal rat kidney (NRK) and mouse embryonic stem cells (mESCs). In Cao et al.^59^, treatment of HeLa cells with siRNA for the SPIN90 activator significantly decreased the actin cortex thickness while concomitantly increasing the cortex density. In these two studies, an increase of actin density with decreasing cortex thickness was thus observed. In mice oocytes, the cortex is a few micrometers thick, which allows for direct optical thickness measurement. Bulteau et al.^60^ showed that the cortex of oocytes obtained from aged mice was thicker and less intense in actin labeling, comparatively to the one of young mice. This change was also correlated to a trend of softer elastic modulus for the thicker cortices. In our study, we quantitatively show that the scaling between thickness and elastic modulus, explained by a constant actin content, is a general property of actin cortices, observed at the single cell and population levels and across a variety of cell types.

### The Dynamic Nature of the Cortex

A crucial difference between cellular solids material such as wood or porous bone and the actin cortex is that the former is dynamic at a time scale of 10s of seconds. By increasing and decreasing the thickness of the actin network with a nearly constant quantity of filamentous actin, the cell is thus able to change drastically its relative density. Perturbations of cytoskeleton regulation pathways such as ROCK and LIMK indicate that the cell is able to modify its thickness and thus its density which have a major effect on its material property (see Fig. 2B). The ability to dynamically shift a material’s position within an Ashby plot distinguishes the actin cortex from virtually all conventional engineered materials. Before discussing the putative physiological functions of this ability, we will discuss the consequences of the discovered scaling on cell mechanics and the other implications of the dynamic nature of the cortex.

On time scales larger than its renewal time, the cortex is considered as a viscous fluid in active matter theoretical models^61,62^. In our experiments, when force is initially relaxed from a limited to a negligible force prior to compression, the cortex thickness increases over a few seconds (see Fig. 1 and Supp Fig. 5). This may be partly due to a viscoelastic relaxation of the material to an unstressed state, but it could also arise from a dependence of the thickness to the applied force, since thickness depends on polymerization speed, which itself could depend on the applied stress. However, at the short timescale of our compression application (1s), actin turnover is expected to be limited, and, accordingly, we observed limited viscous effects during compressions: the temporal delay measured between force and deformation (Supp. Fig. 7B- C) is variable (standard deviation 213 ms) probably because of fluctuations in the cortex thickness^45^ or noise in the distance measurement. The median value is very close to 0, but slightly negative (− 30 ms), which may be due to a limited viscous component acting in parallel to the dominant elastic part.

After each phase of compression-relaxation, the cortex thickness recovered to a value close to the initial one, suggesting an elastic-like behavior on the second timescale. It can however slightly differ from the initial value mainly as a result of the indentation-independent thickness- fluctuations due to myosin II activity as described in Laplaud et al^45^. Nevertheless, during force release, thickness recovery followed a different curve than during the compression phase, thus displaying a clear hysteresis (Supp. Fig. 7A). This hysteresis is a feature already observed on actin networks assembled in vitro^63^. It is often interpreted as a plasticity displayed by amorphous materials that rearrange upon load. Seemingly at odds with this interpretation, repeated measurements at about 19 s interval did not show signs of mechanical fatigue or memory (Supp. Fig. 6). This can be explained by actin turnover in cells, with actin filaments polymerizing and depolymerizing on timescales of tens of seconds, thus resetting the mechanical state between deformations. Understanding this effective almost purely elastoplastic behavior of the dynamic actin network in future studies, using more complex temporal force patterns, will be crucial for interpreting mechanical measurements and predicting how the cortex responds to physiological stresses, such as those encountered during cell migration or division.

### Comparison with AFM measurements

The indentation of living cells by atomic force microscopy has been extensively employed to characterize cellular mechanics^34,38^. These studies use Hertzian contact (or the Sneddon variant for conical indenters) to model the contact geometry and fit the force-thickness curves to obtain a value for the elastic modulus. This approach treats the whole cell as an homogenous material, which is convenient for comparing different situations (malignant vs. normal cells for example^64^) but has obvious flaws: actin cytoskeleton plays a major role in cell mechanics and is extremely inhomogeneously distributed in the cells with high actin density in the cortex and much less in the cytosol. This flaw became apparent when comparing indentations made by different probes^37,65^, which provided, for the same cell type, a value of the elastic modulus that differed by a factor of 10 between sharp cones and large beads^37^. A finite element study of the effect of the indenter shape using a more realistic model with a shell and a cell interior^66^ showed that the cortex (shell) elastic modulus largely dominates the fitted apparent cell elastic modulus for sharp indenters at indentation depths below 400 nm. Despite this precaution, the fitted apparent modulus was still different (by a factor of 2 to 5) from the shell elastic modulus set into the model^66^. This study prompted some subsequent works to use the term “cortical stiffness” for the apparent modulus obtained from shallow indentation using sharp tips^59,67^. Moduli obtained from spherical indenters or without indentation limit were by contrast called “cell stiffness” ^67^. Although they have the same dimension, the material elastic modulus and the stiffness are not the same physical quantities, as the second is not an intensive property of the material and depends on its geometry. Indeed, the apparent modulus fitted from finite element studies was found to increase critically when the model cortex thickness was increased in both the case of sharp and round tips^66^.

In this study, we evidence a strong dependence of the cortex elastic modulus on its thickness, via its density. This coupling could mitigate the impact of cortical thickness on the apparent moduli measured through sharp indenters. Indeed, a thicker cortex should increase the apparent elastic modulus^66^, but also lower the intensive elastic modulus by the modulus-density scaling discovered here. These effects might partly cancel each other, explaining the reduced range of apparent cortex stiffness measures produced with a sharp AFM tip, compared to the range of cortex modulus we observed in a population of cells. It also justifies the use of the sharp indenter AFM method for comparative studies. The cortex elastic modulus we measured with the cell pincher is in the upper range or slightly higher than the apparent elastic modulus measured in the literature (1-13 kPa) which is expected from the above reasoning^37^.

### Different cortex deformation modes

As a thin elastic structure, the cortex can deform through multiple modes: local indentation, bending, and stretching (see Table 1). The force needed to deform the material through these modes depend differently on its thickness *h* and its elastic modulus *E*. For example, in a local indentation of depth *d* (where *d* ≪ *h*) the force depends only on *E* and not on *h.* For larger indentations however, depending on the boundary conditions, the sheet might deform by stretching or by bending. The force needed to stretch the sheet and increase its area is proportional to the *area expansion modulus Ka*, which increases linearly with thickness (*Ka ∼ E h*) in a linear material. To bend a thin sheet in the transverse direction the force is proportional to the *bending modulus B*, which strongly depends on the sheet thickness (*B ∼E h*^3^).

**Table 1.**
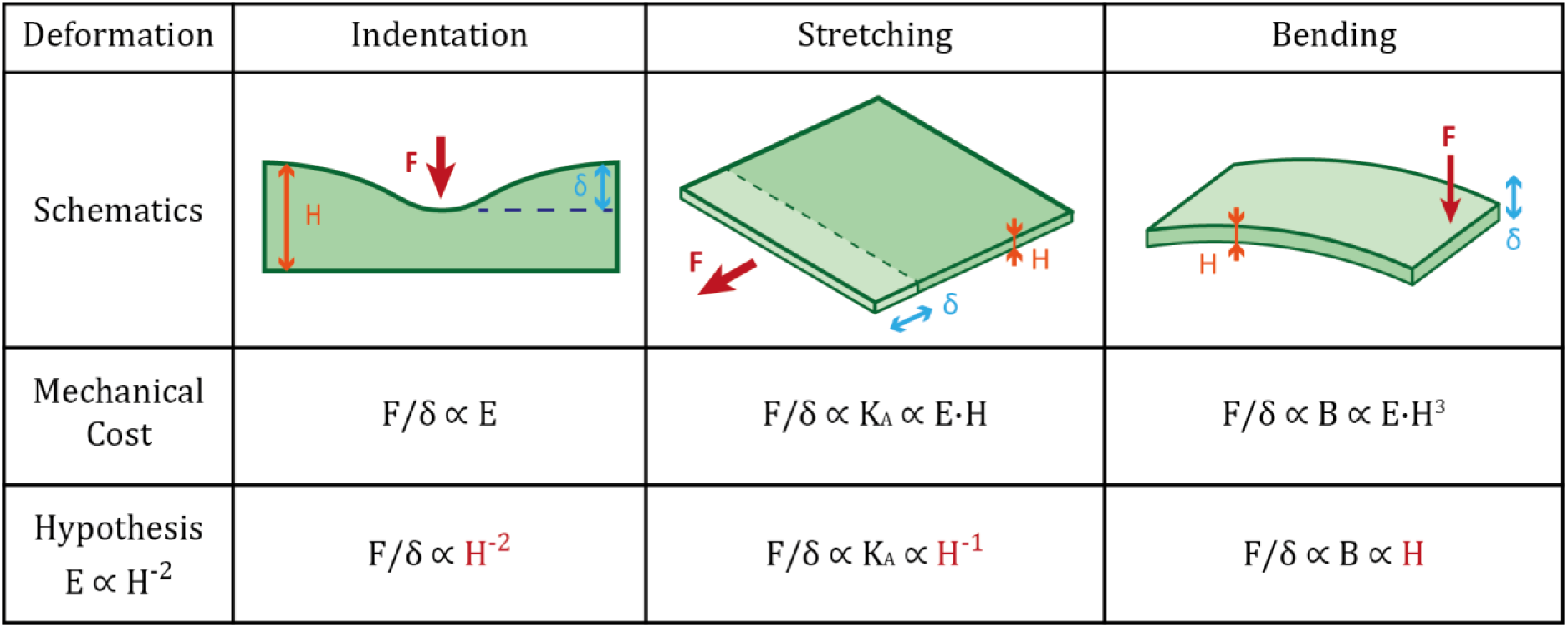
Mechanical costs of different modes of deformation for the cell cortex. . The mechanical cost we define here is the force necessary to deform the material over a given length. In the expressions, K_A_ refers to an area expansion modulus and B to a bending modulus.

| Deformation | Indentation | Stretching | Bending |
| --- | --- | --- | --- |
| Schematics |  |  |  |
| Mechanical Cost | $F/\delta \propto E$ | $F/\delta \propto K_A \propto E \cdot H$ | $F/\delta \propto B \propto E \cdot H^3$ |
| Hypothesis<br>$E \propto H^{-2}$ | $F/\delta \propto H^{-2}$ | $F/\delta \propto K_A \propto H^{-1}$ | $F/\delta \propto B \propto H$ |

The scaling between elastic modulus and thickness that we discovered (*E∼h^-^*^2^) has strong consequences on how these two modes of deformation depend on the thickness. In the case of stretching the dependence of the force on thickness is inverted (*Ka ∼h^-^*^1^) meaning that a thin cortex is harder to stretch than a thick one, contrary to a sheet made of a material whose thickness varies at a constant density. In the case of bending, the impact of increasing the thickness is strongly mitigated (*B ∼ h*). These new relations should hold true in general for any material that changes dimension while maintaining a constant amount of components and thus gets diluted when it increases in size - a phenomenon that has been proposed to generate super-elastic behaviors of cell monolayers^68^. The above analysis helps compare our results to other methods and further explore the consequences of the stiffness-thickness coupling.

In AFM experiments with sharp tips, indentations of a magnitude close to the cortex thickness (400 nm), will result in a mechanical response dominated by indentation (*E ∼ h^-^*^2^), with a small contribution of stretching (*Ka ∼ h^-^*^1^) and bending (*B ∼ h*). In contrast, larger indentations or indentations with a larger tip are expected to increase the contribution of stretching (due to the cell conserved volume at short time scales^69^). Given the different dependence on *h* of these modes of deformation, it is hard to predict the impact of cortex thickness on these measures. Conversely, global measures made with a wedged AFM cantilever on non-adherent cells^43,44^ should depend only on the expansion modulus of the cell cortex, as the cortex expands when the cell is flattened. In these experiments, the initial prestress due to cortex active tension (Myosin II activity) dominates the mechanics, and precise imaging of the cell shape or accurate modeling of the cell geometry is needed to extract an expansion modulus. Different studies report vastly different values for *Ka* in interphase cells, from 1 mN/m ^44^ to several N/m^43^ . This corresponds to a variation of the cortex elastic modulus from ∼kPa to ∼Mpa, our measurements naturally falling between these bounds. Further experiments combining a magnetic pincher and a wedged AFM cantilever would allow a direct verification of the relation *Ka ∼h^-^*^1^ at the single cell level.

### Consequences of the modulus-thickness scaling for cell and tissue morphogenesis

Beyond the comparison with other methods to measure cellular mechanical properties, how different modes of deformation depend on cortex thickness may have major consequences to understand how cells can control their shape both autonomously and in response to external forces. By increasing, locally or globally, the thickness of the cortex, the cell surface can become easier to indent or easier to stretch, but more difficult to bend. Inversely, by thinning and densifying its cortex, the cell surface would become easier to bend but more difficult to indent or stretch. A thicker cortex, by favoring indentations, might facilitate adhesion to the nanoscale topography of the extracellular matrix, or initiation of phagocytosis of small particles. Inversely, a thin cortex, easier to bend, would favor the formation of large folds and ruffles on the cell surface. These structures could then be stabilized by thickening the cortex. Modulation of cortex properties to generate asymmetric daughter cells during division has already been proposed^70^. The difference in modes of deformation due to the modulus-thickness scaling could provide an alternative physical explanation for this phenomenon: considering two daughter cells with different cortex thicknesses, the daughter cell with a thicker cortex should be larger because easier to stretch under a similar internal pressure.

At the tissue level, the mechanical properties of individual cells contribute to collective behaviors, such as tissue folding, wound healing, and embryonic development. The ability of cells to fine-tune their cortex mechanics through thickness modulation could enable coordinated shape changes and force generation across cell populations. Future studies could explore how variations in cortex mechanics influence tissue-level processes, using model systems such as epithelial sheets or organoids combined with the magnetic pincher.

### Conserved actin quantity in cell cortices at the origin of the modulus-thickness scaling

The conservation of the modulus-thickness power-law scaling across different cell types and organisms, with an exponent close to -2, is a striking discovery that suggests strong evolutionary constraints on the structural organization of the actin cortex. Our interpretation of this exponent from a simple scaling reasoning, combined with experimental evidence, led us to conclude that the coupling is the result of a limited variation in the amount of actin in the cortex of a cell not only over time (40%, Supp. Fig. 16 B), but also between cells in a population (66%, Supp. Fig. 16 A) and even between cell types (20%, when comparing central tendencies of cell types, see Supp. Fig. 16 C). This conservation of the amount of cortical actin would mean that the cortex thickness constitutes the main physical ‘knob’ that cells possess to modulate their cortex properties - since thickness changes will lead to density changes and thus to stiffness changes.

Whether the cell modulates cortex thickness through direct physical swelling or dynamical turnover is still an open question. Previous studies have shown that faster polymerization, which should produce thicker cortices, also leads to lower actin density^33,71^. A change in the ratio of nucleation of new filaments *versus* actin disassembly could also dynamically create cortices of variable densities. The limited but significant effect on cortical thickness we observed by perturbing myosin and cofilin points towards an integrated role of numerous cortex effectors. The ubiquity of the modulus-thickness scaling raises the question of the evolutionary constraint that defines the amount of actin filaments in the cortex, and how this constraint imposes this specific mechanism for cell and tissue morphogenesis.

## Conclusion: A Mechanical Framework for Cell Shape and Function

In summary, our study reveals that the actin cortex operates within a constrained mechanical regime, where thickness and elastic modulus are inversely coupled through variations in actin density, as is the case for open cellular solids. This relationship, robust across cell types and perturbations, provides a unifying principle for understanding how cells regulate their mechanical properties and their shape. By modulating cortex thickness with constant actin quantity, cells can fine-tune their stiffness to meet the demands of morphogenesis, migration, and division. These insights not only advance our understanding of cell mechanics but also offer a foundation for exploring how mechanical properties of the actin cortex shape tissue dynamics and organismal development.

