## Supplementary Materials for "Softer when Thicker: a Conserved Scaling Law Governs Actin Cortex Mechanics"

#### *Supplementary Figures*

**Authors**

Joseph Vermeil[1,2], Anumita Jawahar[1,2], Valentin Laplaud[1,2], Eloise Halouchery[1,2], Hugo Lachuer[3], Camille Plancke[2], Laura Bernard[2], Nicolas Borghi[3], Matthieu Piel[2]*, Olivia du Roure[1]*, Julien Heuvingh[1]*

**Affiliations**

1. Physique et Mécanique des Milieux Hétérogènes, CNRS, ESPCI Paris, Université PSL, Sorbonne Université, Université Paris Cité, Paris, France

2. Institut Curie and Institut Pierre Gilles de Gennes, PSL University, CNRS, Paris, France.

3. Institut Jacques Monod, CNRS, Université Paris Cité, Paris, France

*

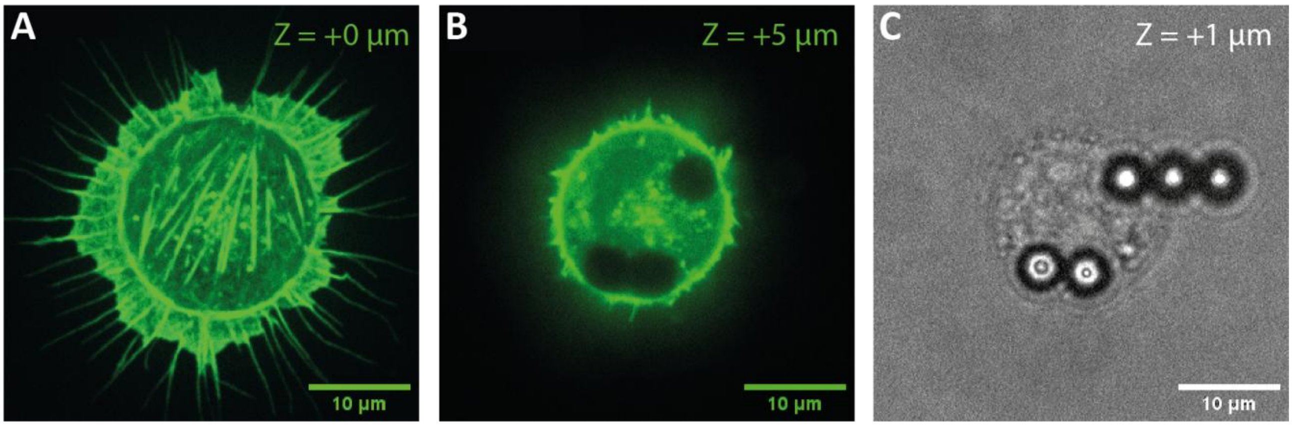


**Supplementary Figure 1 – Pinched cortex of a 3T3 cell on a fibronectin disc**
One 3T3 fibroblast adhering on a 20 um fibronectin disc and prepared for the Magnetic Pincher, with LifeAct-EGFP labeling the F-Actin (green). **A** – The basal plane of the cell. **B** – The plane containing the beads. **C** – Bright field image of the same cell.
100X magnification, scale bars 10 um

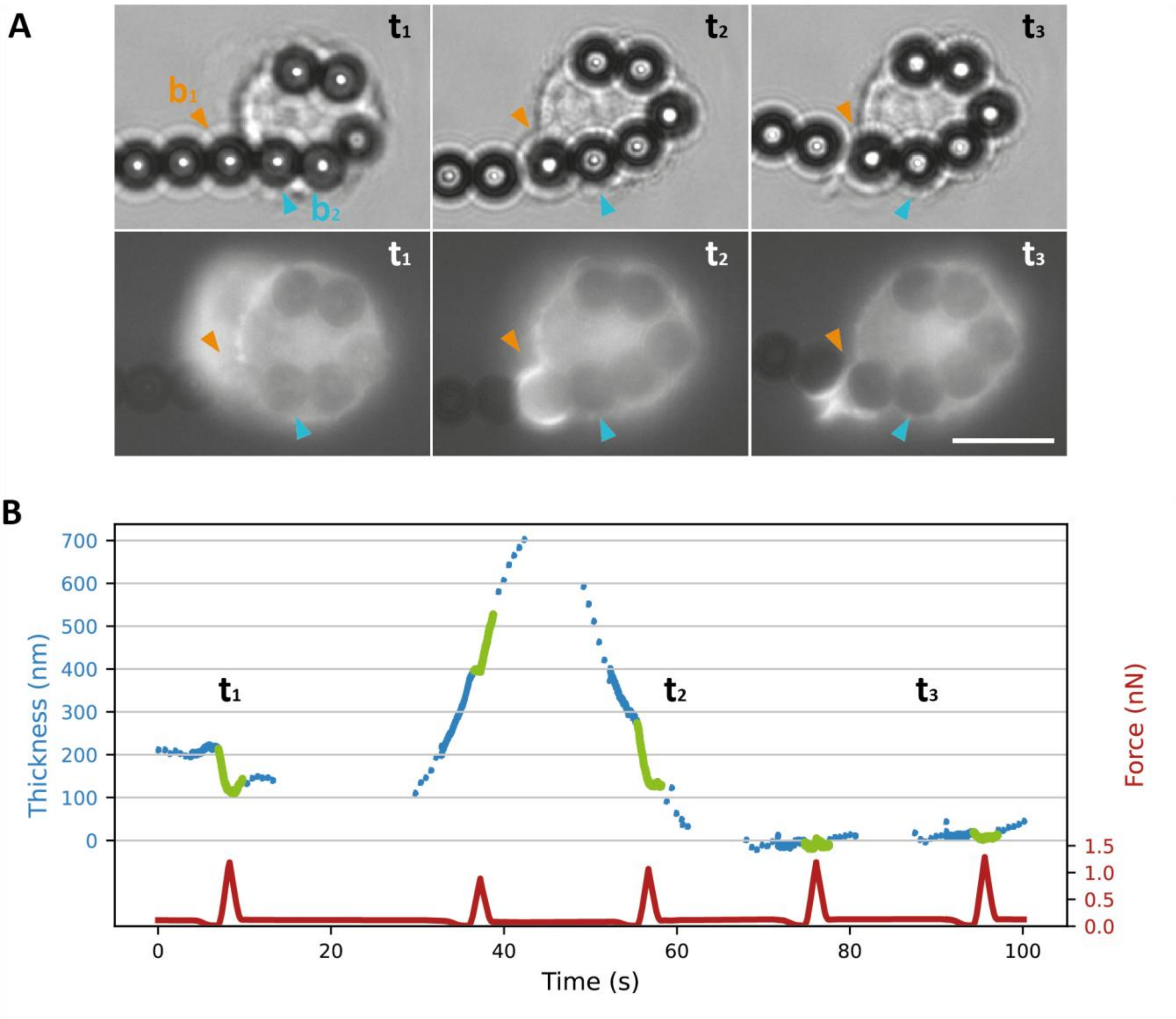


**Supplementary Figure 2 – Beads are internalized quickly and do not seem to retain actin structures inside the cell.**

**A –** Process of bead internalization during a Magnetic Pincher experiment on HoxB8 macrophages, prone to internalize beads. The sequence is shown in bright field (top) and epifluorescence (bottom, LifeAct). The bead b1, outside at t = t1, gets covered by an actin cap by t = t2 and internalized by t = t3. Scale bar 10 um.

**B –** Distance between the beads b1 and b2. At t1, there is still a 200 nm thick layer between the beads, which can be indented. After a transient increase of thickness, the distance between the beads become roughly zero, and is not decreased by peaks of forces.

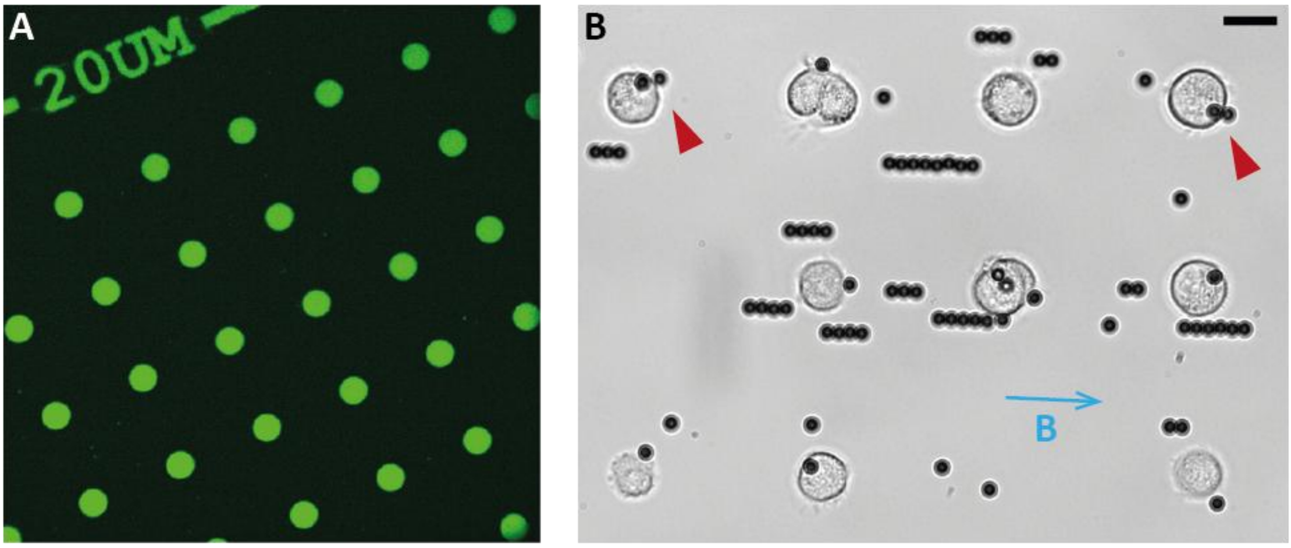


**Supplementary Figure 3 – Fibronectin micropatterns allow for arrays of cells in similar geometries, simultaneously pinched by magnetic beads.**
**A –** Micropatterned discs of fibronectin on a PLL-g-Peg coated glass substrate. The discs are visualized thanks to fluorescent fibrinogen which was added to the fibronectin solution. The discs are 20 um in diameter, and distant by 70 um (center to center).
**B –** 3T3 fibroblasts prepared for a Magnetic Pincher experiment, and adhering on fibronectin discs. Beads attract each other upon exposition to the magnetic field (blue arrow). The two orange arrows show cells which are properly pinched by a pair of beads. 40X magnification, scale bar 20 um.

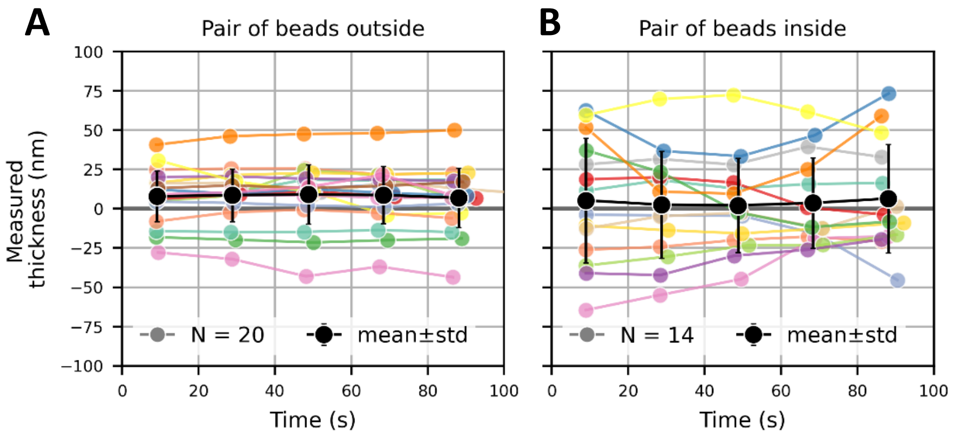


**Supplementary Figure 4 – Precision and variability in bead-to-bead distance measurement.** Distance between two beads forming a pair (surface to surface distance) as a function of time, when the external magnetic field applied on the beads if set at 5 mT (resting value). A – Both beads are outside cells. B – Both beads are inside a cell. In both graphs the black points are the average distance with error bars showing the standard deviation.

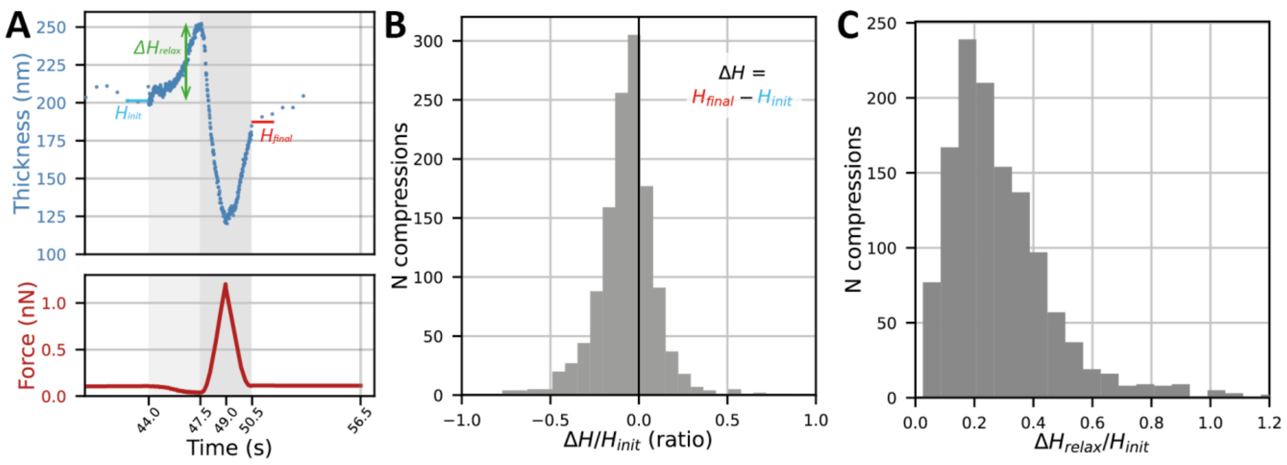


**Supplementary Figure 5 – Typical effects of the relaxation-compression-relaxation scheme on the cortex thickness**

**A –** One example compression with the notations used for panels B and C. $H_{init}$ is the thickness under resting field (5 mT) just before the initial relaxation (cyan mark), $H_{final}$ is the same but just after the final relaxation (red mark) and $\Delta H_{relax}$ is the increase of thickness during the initial relaxation when the field goes from 5 to 2 mT (green arrow), corresponding typically to a force going from 50-70 pN to 15-20 pN.

**B –** Distribution of the ratio $\Delta H/H_{init}$ where $\Delta H$ is defined as $H_{final} - H_{init}$, for a large dataset of compressions on the cortex of DMSO treated 3T3 fibroblasts.

The median value is -0.058. This indicates that the compression sequence has little lasting effects on the cortex, since its final state is almost the same than its initial state, in average. n = 1265 compressions.

**C –** Distribution of the ratio $\Delta H_{relax}/H_{init}$, for the same dataset as panel B.

The median value is 0.25, and there is no accumulation of values close to zero. This indicates that despite the repeated compressions, the cortex always increases its thickness as a response to a small decrease of applied force, which can be seen as elastic relaxation or active reorganization. n = 1265 compressions.

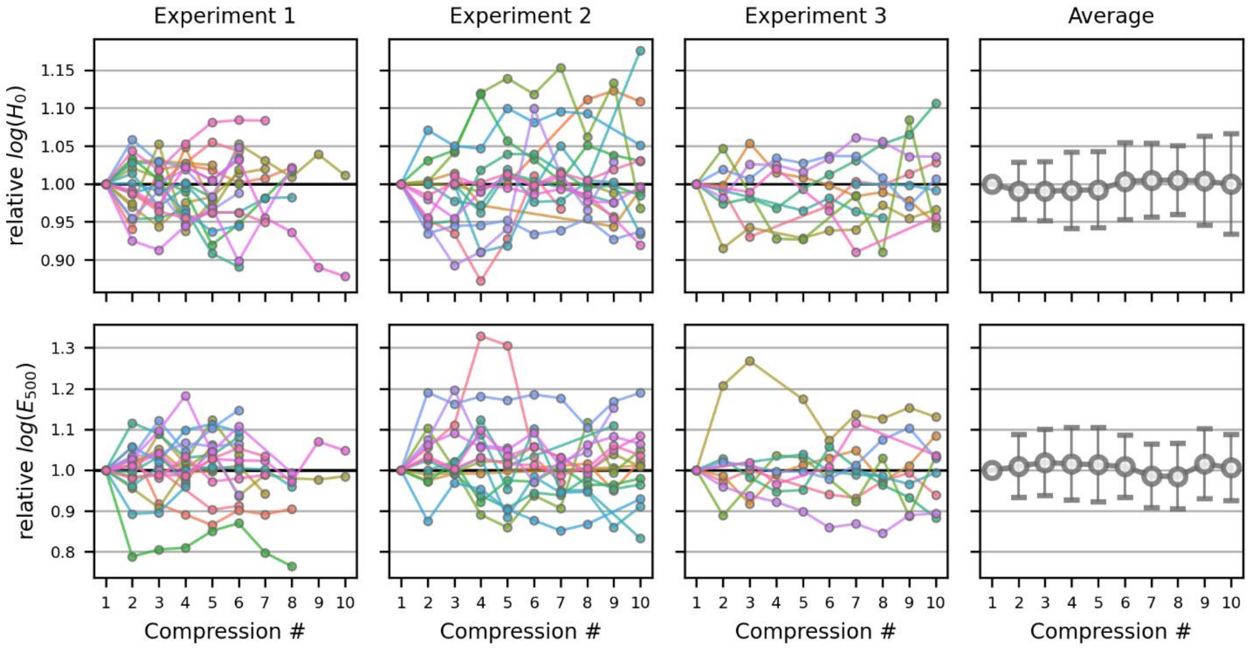


**Supplementary Figure 6 – Cortices show no significant response to repeted indentations.**
For three sets of measurements corresponding to three independant experiments (columns 1-3) we computed the log of the thickness $H_0$ and the elastic modulus E, normalized by the value obtained for each cell's first cortex indentation. Then we plotted the evolution of this relative log-thickness (top row) and log-modulus (bottom row) as a function of repeated compressions of the cortex. The goal was to assess whether cortex mechanics would be shifted in a typical way as compressions are repeated every ~19 seconds. The last column shows the average of all cells of experiments 1, 2 and 3. Since the average relative log-thickness and the average relative log-modulus are both stable around 1 when compressions are iterated, we conclude that in the precise conditions of our measurements, the repetitions of cortical indentations does not affect the cortex thickness and elastic modulus.
All three experiments were performed on DMSO treated 3T3 fibroblasts.
Expt 1: n = 122 compressions on N= 21 cells.
Expt 2: n = 75, N = 20.
Expt 3: n = 377, N = 29.

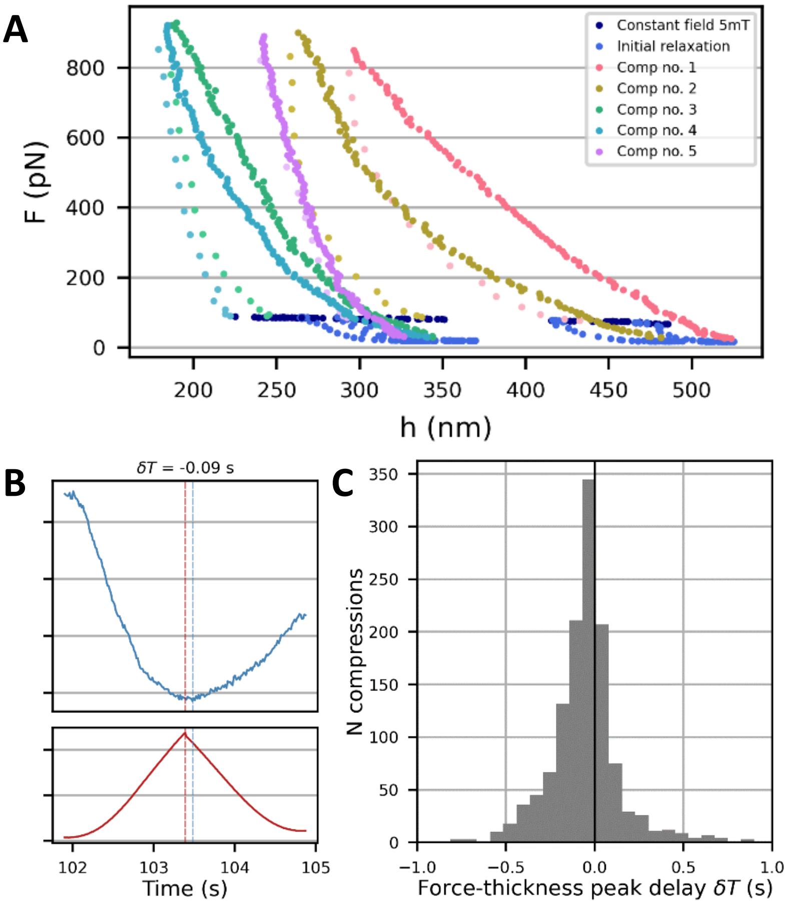


**Supplementary Figure 7 – The distribution of delays between the peak of force and the thickness minimum points toward a minor contribution of the effective cortical viscosity.**

**A –** Series of compressions on the same cortex. Each of the 5 cycles is represented in a different color, with the points corresponding to the indentation in a darker shade than those corresponding to the compression. In blue are the points corresponding to the initial relaxation before each indentation, and in darker blue are the points corresponding to the phases where the field is maintained to its resting value (5 mT).

**B –** Example of delay determination in one typical compression. The red dashed line shows the time of the force maximum ($t_{FM}$) and the dashed blue line shows the one of the thickness minimum ($t_{Hm}$). In that case $\delta T = t_{FM} - t_{Hm} = -0.09$ s.

**C –** Distribution of delays δT for a large dataset of compressions on the cortex of DMSO treated 3T3 fibroblasts. The median value is – 0.03 s, which suggests that the viscous contribution to the cortex mechanics is small for these compressions. n = 1265 compressions.

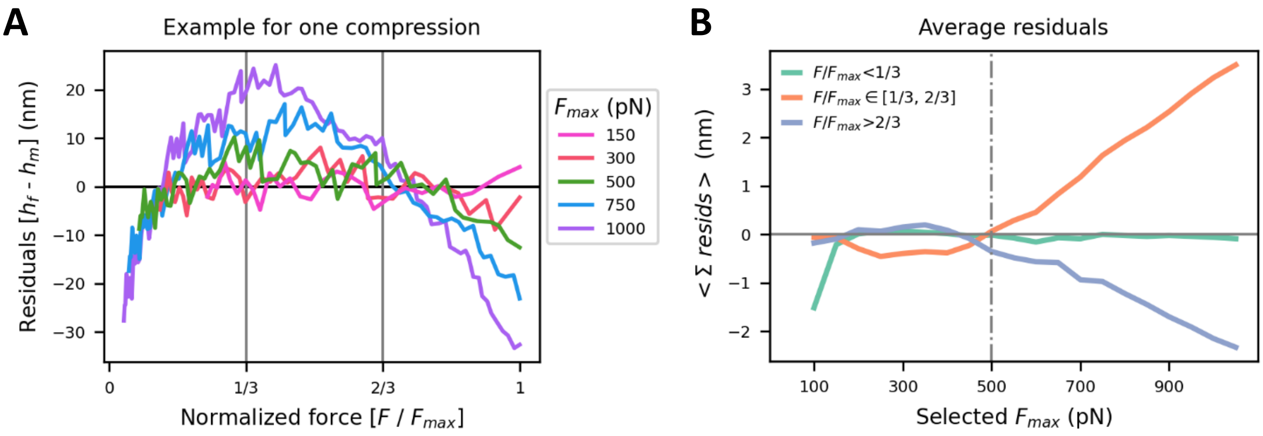


**Supplementary Figure 8 – Choosing the upper bound $F_{max}$ = 500 pN in force-thickness fits.** We chose to fit over the widest range of force [0, $F_{max}$] that would keep the residuals small and unstructured. In the force-thickness curves measured from cortex indentations, we considered the force F as the least noisy quantity (it is mostly controlled by the external magnetic field that we set) compared to the cortex thickness h (which is only measured). Hence we fitted our force-indentation model with force as the X variable and thickness as the Y variable.

**A** – Residuals ($h_{fit}$ – $h_{measured}$) of the Chadwick model fit as a function of the force, with diverse values for the upper bound of the fit $F_{max}$, in the case of one typical compression. In order to analyse the structure of the residuals in a simple way, we normalized the force by Fmax and split the range of normalized force in three regions, from 0 to 1/3, from 1/3 to 2/3 and from 2/3 to 1. On this typical compression we see that for Fmax ≤ 500 pN the residuals are relatively unstructured, while for higher values of the upper bound $F_{max}$, they become distinctively structured (high in the second third, low in the last third).

**B** – Average sum of residuals for a large dataset of compressions on cortices of DMSO treated cells. Each curve represent the average of the sum of residual over one third of the chosen force range, as a function of the selected upper bound Fmax. It confirms the qualitative observation from pannel A: at high values of Fmax, the residuals are clearly structured in average. From this graph, we determined $F_{max}$ = 500 pN (gray dashed line) as the highest value that would keep the residuals unstructured in average.

n = 1539, N = 220, M = 14 (n compressions on N cells over M experiments).

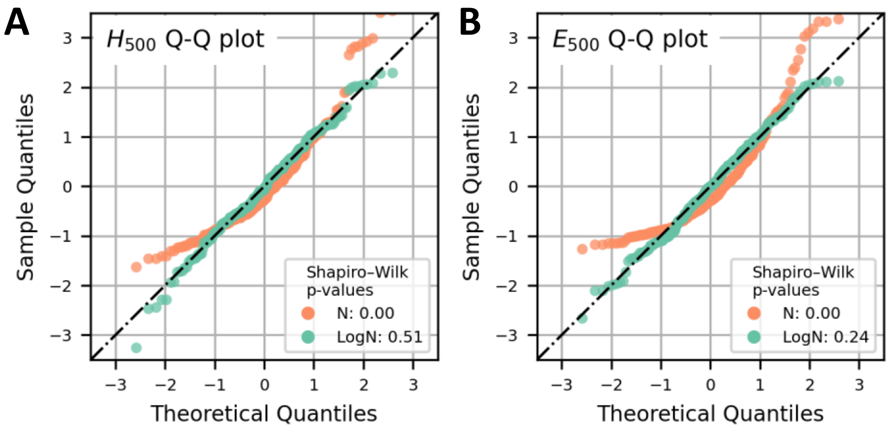


**Supplementary Figure 9 – The measured cortical thickness and elastic moduli follow a log-normal distribution.**

The log-normal nature of these distributions is shown here by quantile-quantile plots (Q-Q plots) for the undeformed thickness (**A**) and the elastic modulus (**B**). Briefly, these represent the percentiles of a normalized series of values (substracted by its mean and divided by its standard deviation) versus the expected percentiles of a normal distribution. If the points follow the Y=X line, the considered series of value is normally distributed. Here the orange points show the Q-Q plots for the distributions of $H_0$ and E, while the blue points correspond to the distributions of $\log(H_0)$ and log(E). The fact the blue points are much better aligned with the Y=X dashed line is a qualitative indication that these two distributions are log-normal rather than normal.

As a complement, the legends show the p-values for the Shapiro-Wilk normality test, where the null-hypothesis is that a set of values is normally distributed. This null-hypothesis is rejected for $H_0$ and E, but not for $\log(H_0)$ and log(E), further indication that the two measured quantities are log-normally distributed.

As a consequence of this finding, the means and standard deviations of $H_0$ and E are never computed in this study. Instead, we considered the mean and standard deviations of the log of those two quantities.

n = 1265 compressions.

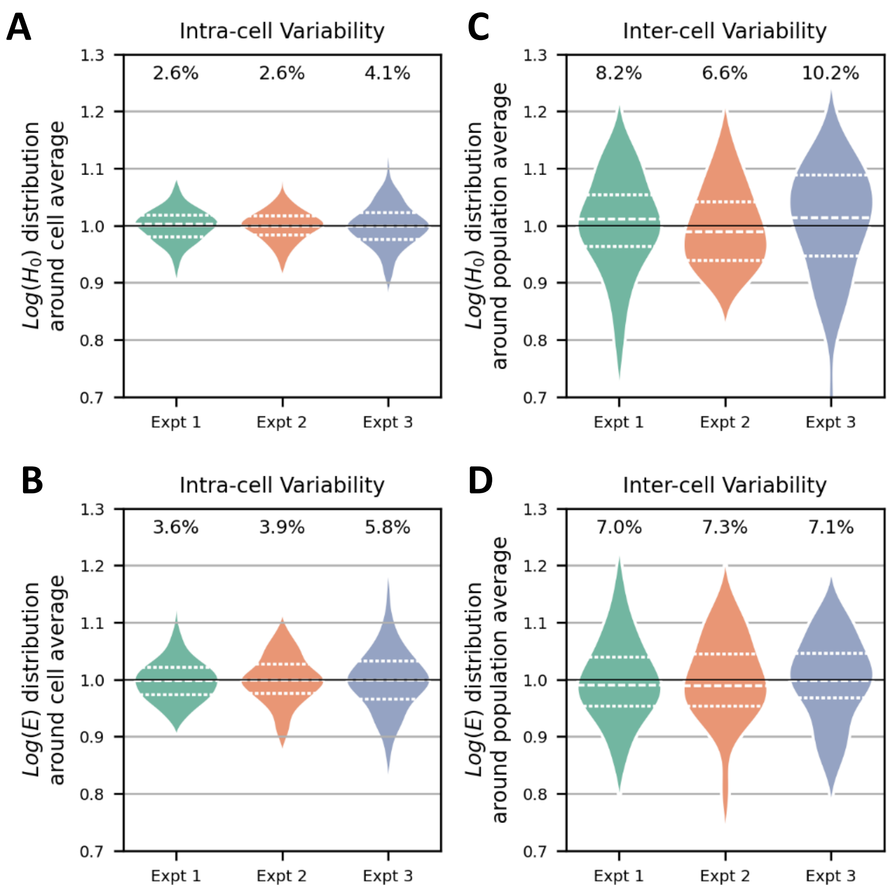


**Supplementary Figure 10 – Intra-cellular versus inter-cellular variability**

For both undeformed thickness H0 and elastic modulus E we computed the intra-cellular variability (ie how measures vary in time for one given cell) and the inter-cellular variability (ie how measures vary from cell to cell within a population). This was done on three sets of measures corresponding to three typical experiments (Expt 1, 2 and 3 in the graphs). Given the log-normal nature of the distributions for $H_0$ and E, we proceeded as such:

- For intra-cellular variability, we grouped the data by cell and took the log of the values. Then the log-values for each cell were divided by their average. These distribution are shown in **panel A** for $H_0$ and **panel B** for E. The percentages above each distribution indicated the coefficient of variation (CV) defined as mean divided by standard deviation.
- For intra-cellular variability, we took the log of the values. Then the mean of the log-values was computed for each cell, and those were divided by their mean (the population average). These distribution are shown in **panel C** for $H_0$ and **panel D** for E. Again, the percentages above each distribution indicated the CV.

In all violin plots shown here, the white dashed line indicates the median and the white dotted lines show the quartiles.

These graphs are an indication that cells possess a « cortical identity », since the intra-cellular variability is clearly lesser than the inter-cellular variability for both the thickness $H_0$ (2.7-fold in average) and the elastic modulus E (1.6-fold in average).

All three experiments were performed on DMSO treated 3T3 fibroblasts.

Expt 1: n = 122 compressions on N= 21 cells.

Expt 2: n = 75, N = 20.

Expt 3: n = 377, N = 29.

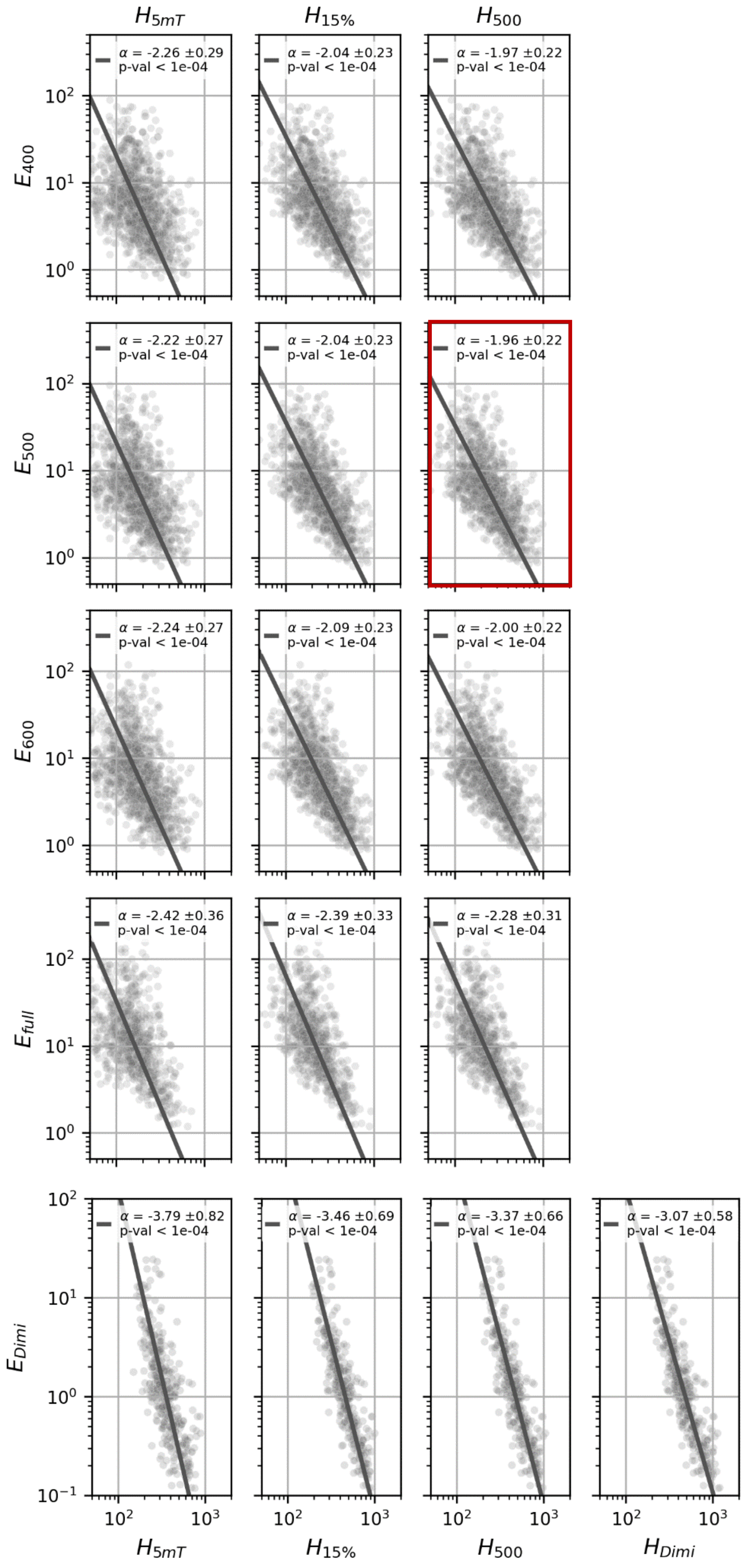


**Supplementary Figure 11 – Thickness-modulus coupling with alternative metrics.**
Legend on the next page

**Supplementary Figure 11 – Thickness-modulus coupling with alternative metrics.**
This grid of graphs show the pair-wise scatterplots of diverse metrics for the cortex elastic modulus (rows) and the cortical thickness (columns), measured on 3T3 fibroblasts in control conditions. The definition of each of these metrics are the following:

- $E_{400}$, $E_{500}$, $E_{600}$ – Elastic moduli fitted with the Chadwick formula on the measured force-indentation curves by setting the upper bound for F as 400, 500 and 600 pN respectively. Note that $E_{500}$ is the metric for the elastic modulus we used througout this study.
- $E_{full}$ – Elastic moduli fitted with the Chadwick formula on the measured force-indentation curves without setting a upper bound for F (fit of the whole curve).
- $H_{5mT}$ – Thickness determined by taking the median of all points between compressions when the applied magnetic field is at its
- $H_{15\%}$ – Undeformed thickness fitted with the Chadwick formula on the measured force-indentation curves by selecting only the points where $F < Fmin + 0.15 \cdot (F_{max} - F_{min})$, i.e. the first 15% of the force range.
- $H_{500}$ – Undeformed thickness fitted with the Chadwick formula on the measured force-indentation curves by setting the upper bound for F as 500 pN. Note that $H_{500}$ is the metric for the elastic modulus we used throughout this study.
- $E_{Dimi}$ and $H_{Dimi}$ – Elastic modulus and undeformed thickness fitted with the Dimitriadis model, only on the region of the force-indentation curves that validated the hypothesis of this model (for more details see Materials and Methods). The curves corresponding to thin cortices were most of the time always outside the model hypotheses, hence these curves were ignored: this is why we show a much reduced number of points for these particular metrics.

All dataset are fitted with a power law $y = A \cdot x^{\alpha}$ using orthogonal distance regression (dark gray lines), α and 95% confidence intervals are indicated in the legends, as well as the Pearson's p-value.
The fact that pair-wise correlations are significant for all metrics demonstrates the robustness of the thickness-modulus coupling.

All data is from DMSO treated 3T3 fibroblasts. Number of replicates for each condition (n compressions, N cells, M experiments) are the following.
For the first 3 rows, n = 1265, N = 210, M = 14
For the $4^{th}$ row ($E_{Full}$), n = 868, N = 180, M = 14
For the $5^{th}$ row ($E_{Dimi}$), n = 414, N = 111, M = 14

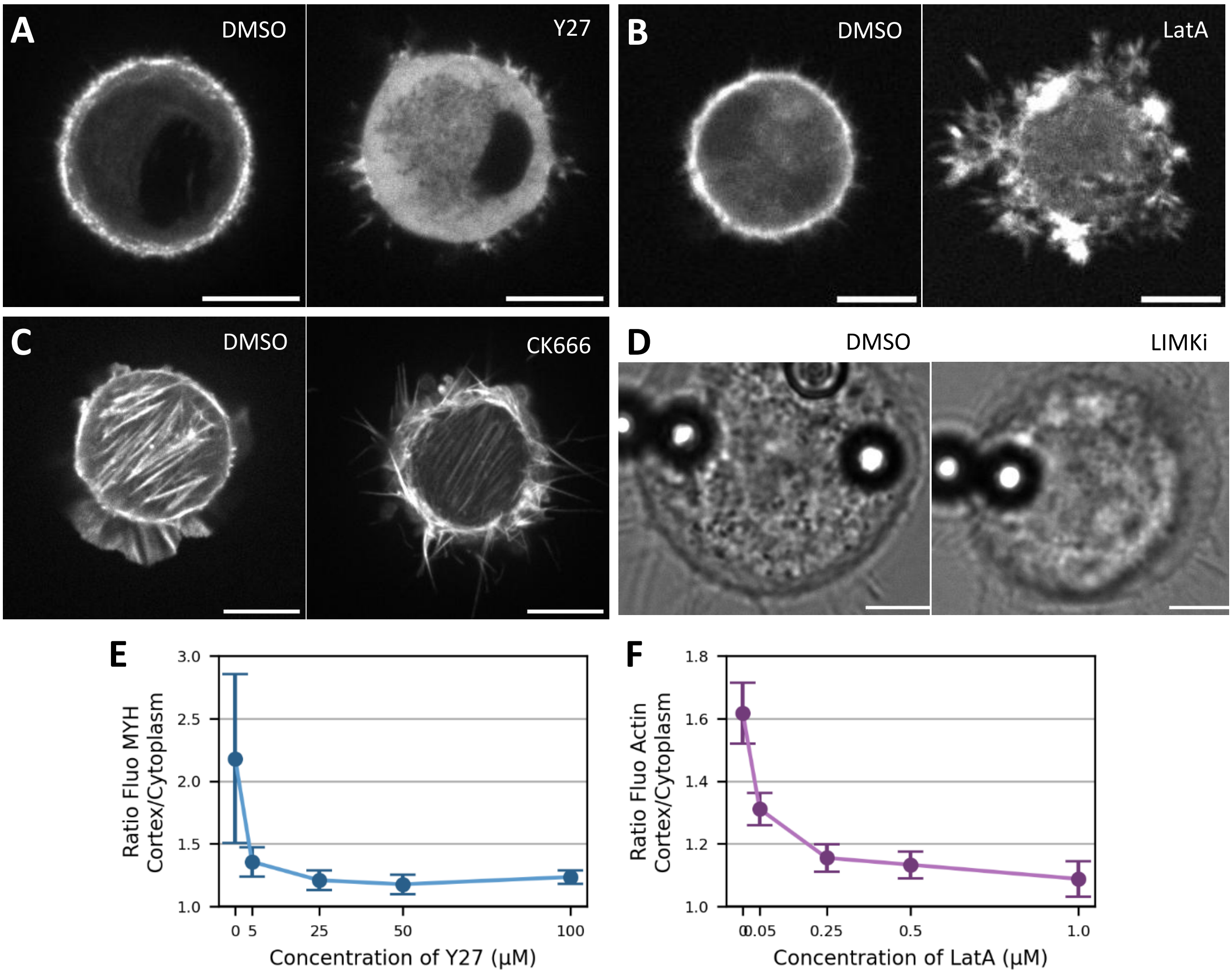


**Supplementary Figure 12 – Drug treatments**

**A-D** – Qualitative effect of the drugs on treated cells, with controls on the left and treeted cells on the right. A – HeLa cells on 20 um fibronectin disc, where the myosin heavy chain (MYH) is tagged with RFP. On the right is a cell treated with 50 µM Y27 (ROCK kinase inhibitor). B – 3T3 fribroblasts on 20 um fibronectin disc, where actin is tagged with LifeAct-EGFP. On the right is a cell treated with 0.5 um LatrunculinA. C – 3T3 fribroblasts on 20 um fibronectin disc, where actin is tagged with LifeAct-EGFP. On the right is a cell treated with 50 uM CK666 (Arp2/3 inhibitor). D – 3T3 fribroblasts on 20 um fibronectin disc with magnetic beads and the external magnetic field turned on. On the right is a cell treated with 20 uM LIMKi (LIMK kinase inhibitor).

**E-F** – Quantification of the dose-dependant effects of Y27 (E) and Latrunculin A (F). Panel E shows the cortex-to-cytoplasm intensity ratio of MYH-RFP in HeLa cells, as a function of the dose of Y27 applied. In the control case the ratio is high, which reflect the fact than in a normal cell the cortex is enriched in myosin 2 compared to the cytoplasm. As the dose of Y27 is increased, the ratio decreases, indicating that myosin 2 is less and less present in the cortex and conversely in the cytoplasm. This is interpreted as a decrease in contractility for the cortex of Y27-treated cells. Panel F is the exact analog, where this time the cells are 3T3 fibroblasts with actin tagged by LifeAct-EGFP. Increased dose of Latrunculin A results in the LifeAct-EGFP tag being more homogeneously distributed in the cells, revealing a loss of filamentous actin at the cortex.

In both cases, the analysed cells were imaged in their middle plane while adhearing on 20 um fibronectin discs. 10 to 25 cells were analysed per concentration. The curves show means ± standard deviations.

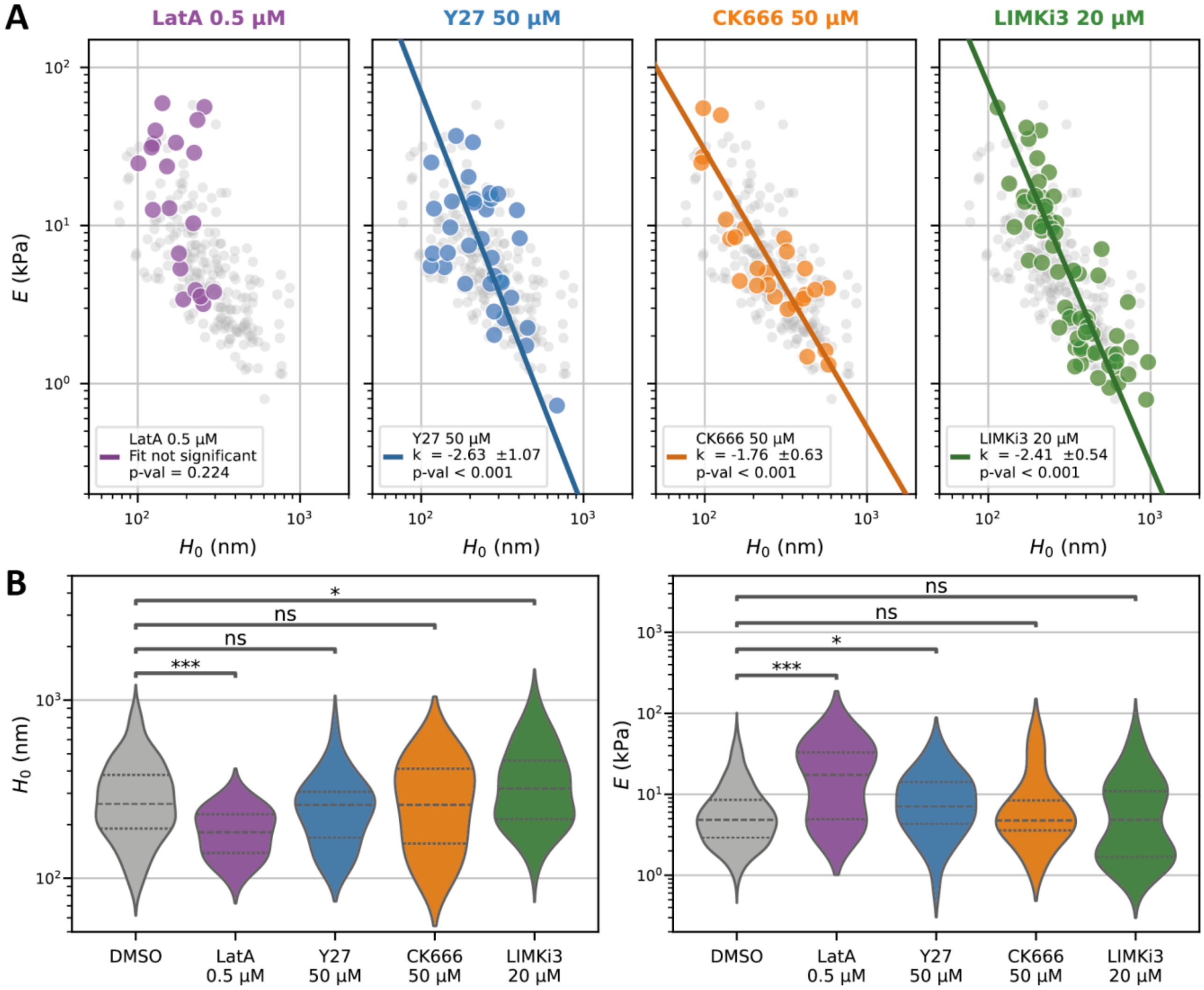


**Supplementary Figure 13 – Drug treatments**

**A** – Effect of diverse drug treatments on the thickness-stiffness coupling. These are the exact same data as Fig. 2B but each point represent the average value for one cell (i.e. the average $H_0$ and E for the series of compressions applied on a given cell). For each treatment the data is fitted with a power law $y = A \cdot x^{\alpha}$ using orthogonal distance regression (colored line), with the data for control cells shown in the background (gray dots). Values of α, 95% confidence intervals and Pearson p-values are indicated on each graph. The fit were shown only if p-val < 0.05.

**B** – Distributions of H0 (left) and E (right) for each drug treatment. Lines inside the violin plots are the median (dashed) and the quartiles (dotted). Lines above show the p-values for the two-sided Mann-Whitney-Wilkoxon non-parametric test (ns: p > 0.05, *: p < 0,05, ***: $p < 10^{-3}$, **** : $p < 10^{-4}$).

See below the number of replicates for each conditions (n compressions, N cells, M experiments).

*DMSO* (control) : n = 1266, N = 210, M = 14.

*LatA* : n = 233, N = 57, M = 3

*Y27* : n = 303, N = 56, M = 3

*CK666* : n = 137, N = 26, M = 2

*LIMKi3* : n = 253, N = 65, M = 3

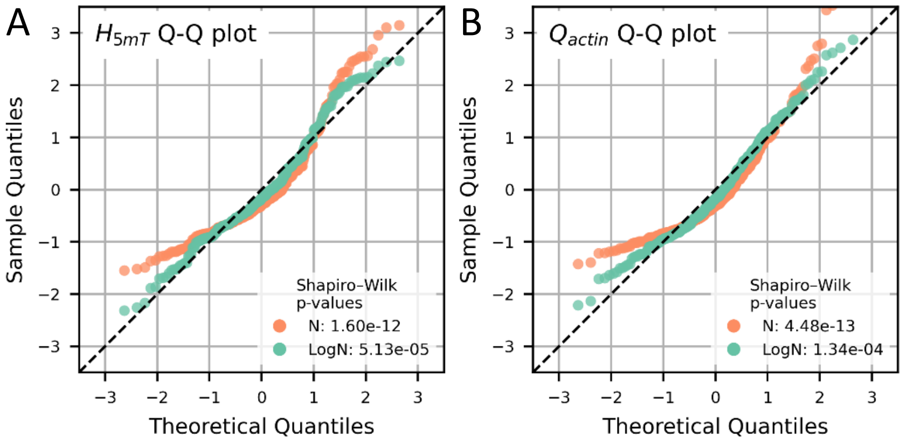


**Supplementary Figure 14 – The measured cortical thickness at resting field and actin quantity at the cortex follow a log-normal distribution.**

The nature of these distributions is examined here by quantile-quantile plots (Q-Q plots) for the thickness at resting field $H_{5mT}$ (A) and the actin quantity at the cortex $Q_{actin}$ (B). Orange points refer to the distribution of $H_{5mT}$ and Q , while blue points refer to the distribution of $\log(H_{5mT})$ and $\log(Q_{actin})$. As a complement, the legends show the p-values for the Shapiro-Wilk normality test. The interpretations of both the Q-Q plots and the Shapiro-Wilk test are explained above in the legend of Supp. Fig. 9.

Here the graph shows that these distribution are clearly not normal, but rather close to be log-normal. The relatively small sample size here might explain the deviation to log-normal distributions. n = 238 points**.**

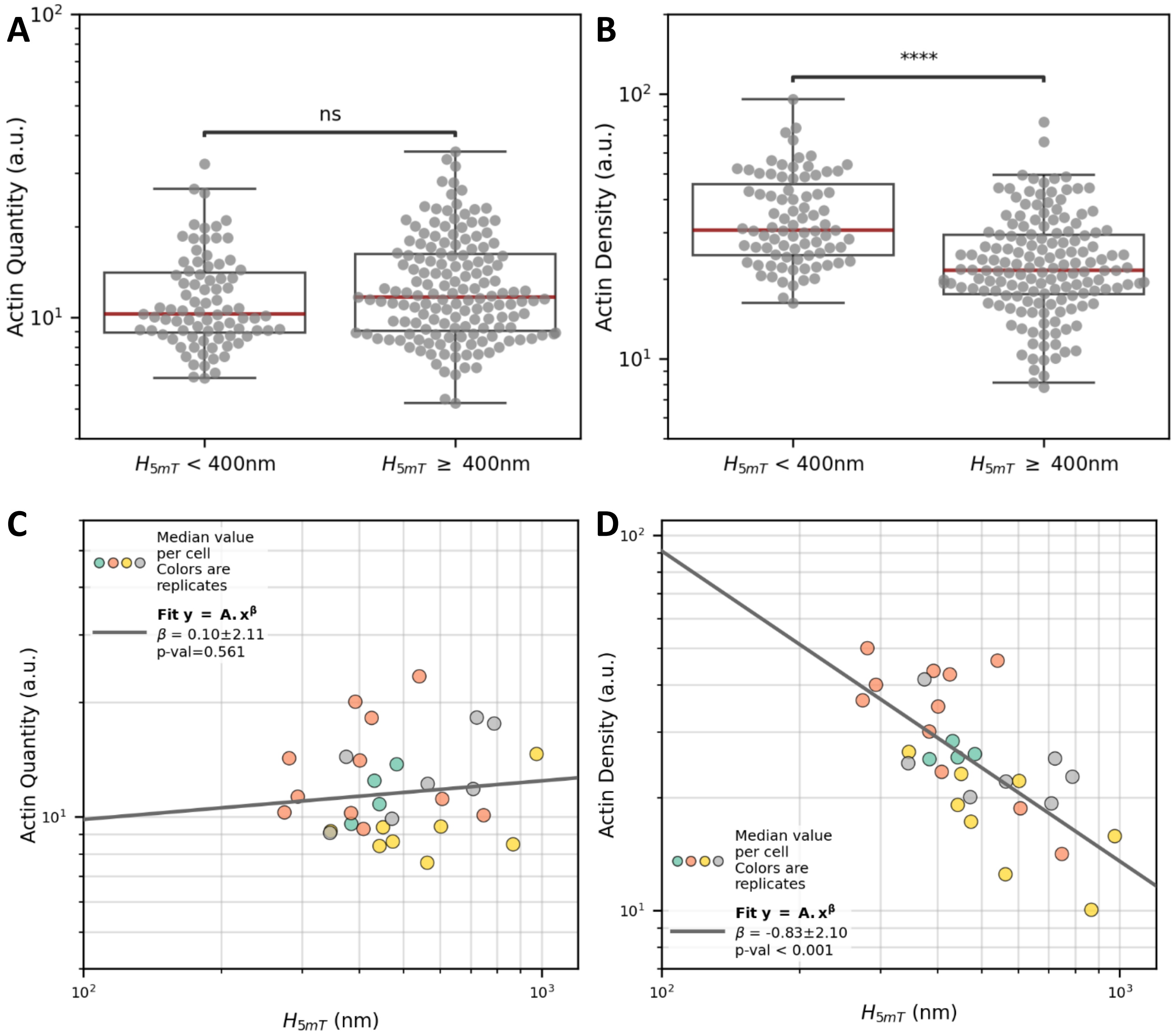


**Supplementary Figure 15 – Cell median actin quantity and density versus median thickness.** A, B – Distributions of actin quantity (A) and density (B) for two bins of thickness ($H_{5mT}$ lower or higher than 400 nm). Like in Fig. 3F and 3G, each point corresponds to a paired measurement of the cortical thickness and the actin quantity at the cortex. Lines above show the p-values for the two-sided Mann-Whitney-Wilkoxon non-parametric test (ns: $p > 0.05$, **** : $p < 10^{-4}$). n = 238 points, N = 30 cells, M = 4 experiments.

C, D – Actin quantity (C) and density (D) versus thickness ($H_{5mT}$) in log-log scale. These graphs represent the same data than Fig. 3F and 3G, except that here each point represents the median of all measures on one cell. Exponents were computed by fitting a power law (robust fit with a Huber cost function, 95% confidence intervals). n = 238 points, N = 30 cells, M = 4 experiments.

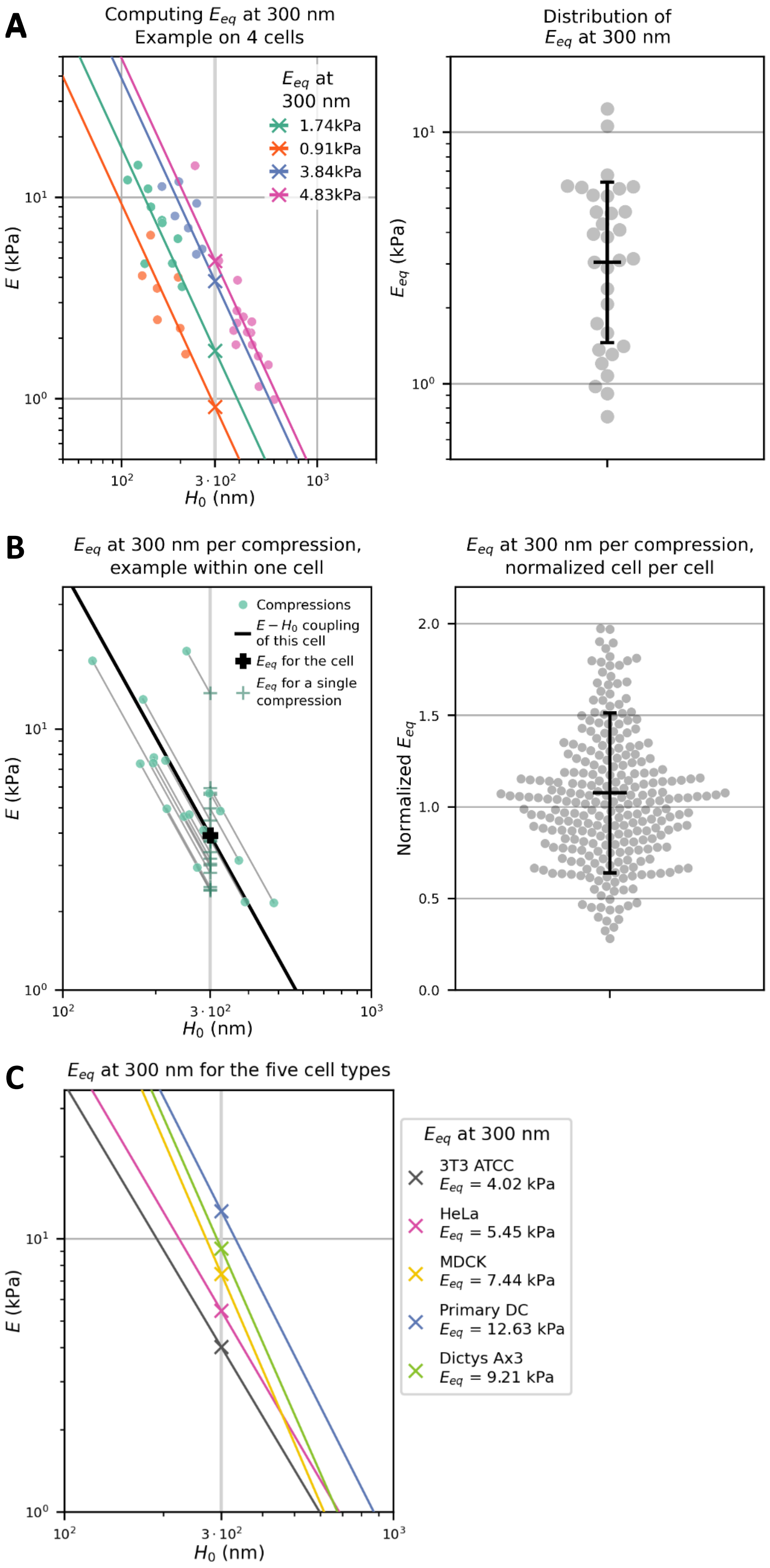


**Supplementary Figure 16 – Comparing power-law scaling using an equivalent elastic modulus at $h_{ref}$ = 300 nm.** See legend on the next page.

**Supplementary Figure 16 – Comparing power-law scaling using an equivalent elastic modulus at $h_{ref}$ = 300 nm.**

**A – Left:** Computation of $E_{eq}$ the equivalent modulus at 300 nm for four cells from the dataset shown in Fig. 4A. The idea is to calculate a value of what would be the elastic modulus of a cortex if it was 300 nm thick, following the thickness-modulus coupling. Briefly, each cell of this dataset was analysed as described in the legend of Fig. 4 in order to determine the exponent of the cell thickness-modulus coupling. The median of these exponents was equal to $\alpha_{med}$ = -2.11 (as seen in the inset of Fig. 4A). Inferring that this value represents well enough the behaviour of all thickness-modulus couplings of this population of cells, for each cell the set of measured E and $H_0$ was fitted with a power-law forcing the exponent to be equal to $\alpha_{med}$ : $y = A \cdot x^{\alpha,med}$. Using these fits, represented on the left for four typical cells, we calculated the modulus $E_{eq}$ defined as $E_{eq} = A \cdot 300^{\alpha,med}$. **Right:** The distibution of all $E_{eq}$ at 300 nm of cells from the dataset shown in Fig. 4A. The black mark shows the log-mean and the log-standard deviation (3.0 kPa, [1.6, 6.3] kPa). The associated coefficient of variation is CV = log-std / log-mean = 66 %. n = 397 compressions, 32 cells, 5 experiments.

**B – Left:** Computation of $E_{eq}$ the equivalent modulus at 300 nm for successive compressions on the same cell. The difference is that now, instead of computing one value of $E_{eq}$ for a cell, we use the same inferred exponent as before ($\alpha_{med}$ = -2.11) to see what each measure of E and $H_0$ would be at 300 nm following the same thickness-modulus coupling. This graph illustrates this calculation with the example of one typical cell, with n = 17 compressions.
**Right:** The distibution of single-compression-$E_{eq}$ at 300 nm of cells from the dataset shown in Fig. 4A. For each cell the single-compression-$E_{eq}$ where normalized by the cell $E_{eq}$, calculated as described above for panel A. The black mark shows the mean and the standard deviation (1.08 ± 0.44). The associated coefficient of variation is CV = std / mean = 40 %. n = 274 compressions, N =17 cells, M = 1 experiment.

**C –** Computation of $E_{eq}$ the equivalent modulus at 300 nm for the thickness-modulus coupling of the five tested cell types. For each cell type, the value of the elastic modulus at 300 nm was infered from the power-law fitted on the E-$H_0$ coupling, seen on Fig. 4B. On this graph only the line representing the fits, and the values of $E_{eq}$ are represented. The coefficient of variation associated with the spread of these values is CV = std / mean = 20 %.

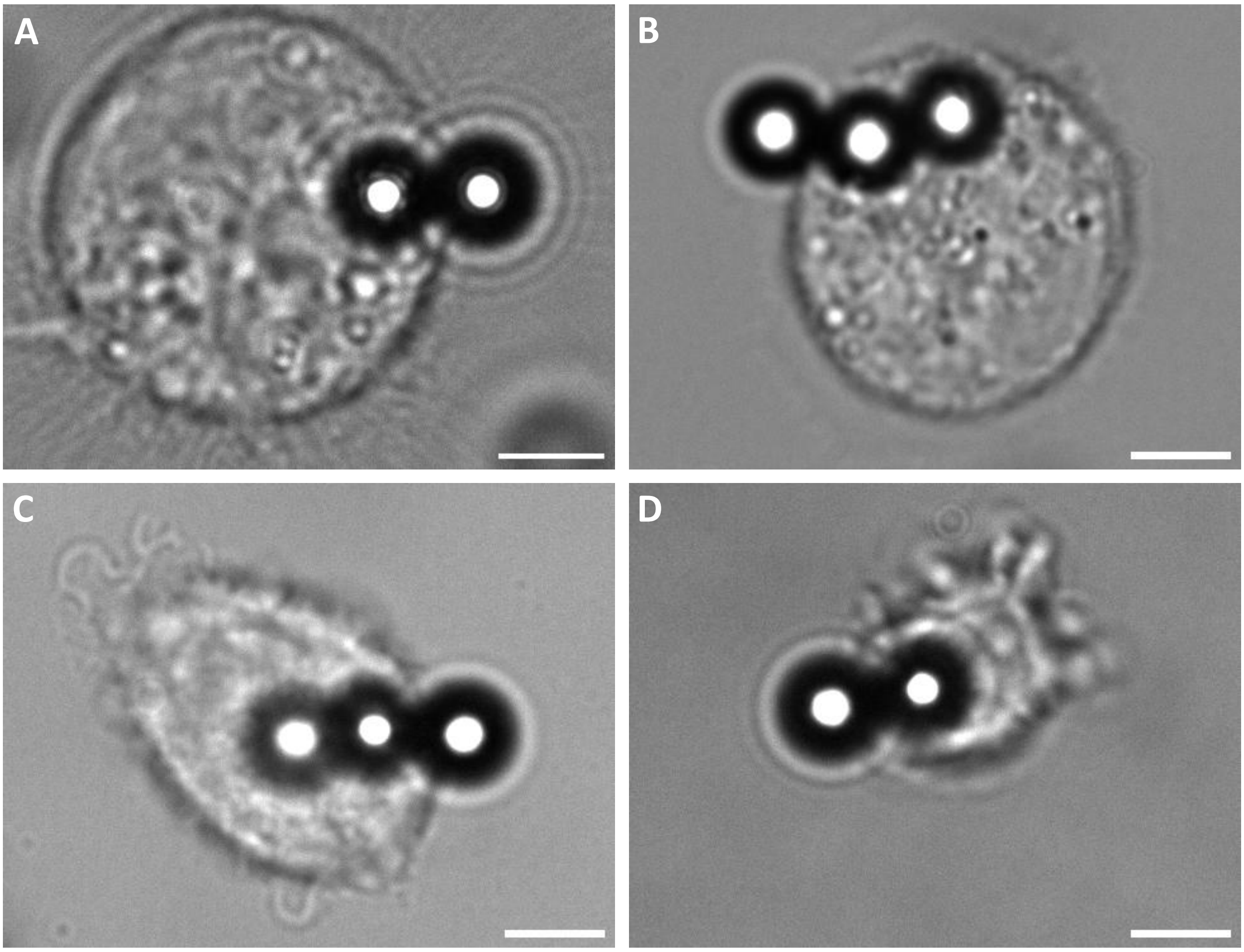


**Supplementary Figure 17 – Aspects of cells from different types when measured with the Magnetic Pincher.**

**A** – HeLa cell on a 20 um fibronectin disc.

**B** – Madin-Darby canine kidney (MDCK) cell on a 20 um fibronectin disc.

**C** – Primary mouse dendritic cell (DC) on a BSA-coated glass surface.

**D** – Dictyostelium Discoideum (Dicty) on a BSA-coated glass surface.

Scale bars 5 um

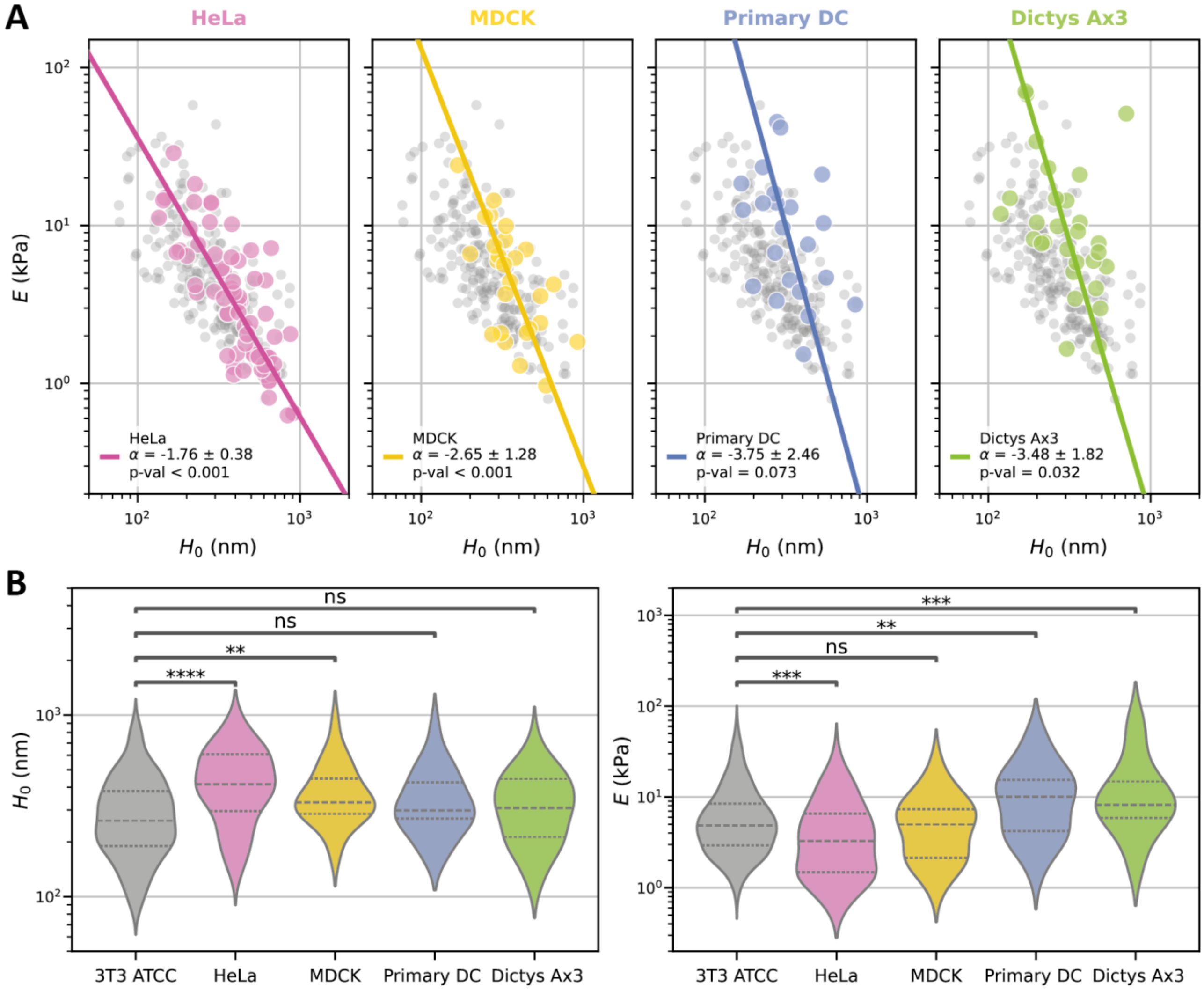


**Supplementary Figure 18 – Cortices of different cell types**

**A** – Modulus-thickness coupling in diverse cell types. These are the exact same data as Fig. 4B but each point represent the average value for one cell (i.e. the average $H_0$ and E for the series of compressions applied on a given cell). For each treatment the data is fitted with a power law $y = A \cdot x^{\alpha}$ using orthogonal distance regression (colored line), with the data for control cells shown in the background (gray dots). Values of α, 95% confidence intervals and Pearson p-values are indicated on each graph.

**B** – Distributions of $H_0$ (left) and E (right) for each cell types. Lines inside the violin plots are the median (dashed) and the quartiles (dotted). Lines above show the p-values for the two-sided Mann-Whitney-Wilkoxon non-parametric test (ns: $p > 0.05$, **: $p < 10^{-2}$, ***: $p < 10^{-3}$, **** : $p < 10^{-4}$).

See below the number of replicates for each conditions (n compressions, N cells, M experiments).

*3T3 ATCC*: n = 1266, N = 210 , M = 14.

*HeLa Fucci cells*: n = 463, N = 67 , M = 5.

*MDCK cells*: n = 128, N = 26 , M = 3.

*Primary DC*: n = 220, N = 22 , M = 6.

*Dictyostelium Ax3*: n = 200, N = 29 , M = 4.

### Softer when Thicker: a Conserved Scaling Law Governs Actin Cortex Mechanics

#### *Materials and Methods*

**Authors**

Joseph Vermeil[1,2], Anumita Jawahar[1,2], Valentin Laplaud[1,2], Eloise Halouchery[1,2], Hugo Lachuer[3], Camille Plancke[2], Laura Bernard[2], Nicolas Borghi[3], Matthieu Piel[2]*, Olivia du Roure[1]*, Julien Heuvingh[1]*

**Affiliations**

1. Physique et Mécanique des Milieux Hétérogènes, CNRS, ESPCI Paris, Université PSL, Sorbonne Université, Université Paris Cité, Paris, France

2. Institut Curie and Institut Pierre Gilles de Gennes, PSL University, CNRS, Paris, France.

3. Institut Jacques Monod, CNRS, Université Paris Cité, Paris, France

*

### I – Materials & Methods

#### 1. Cells & culture

##### i. 3T3 fibroblasts

We purchased cells from the ATCC cell bank from LGC standards (#ATCC-CRL-1658) in January 2023. They were amplified and frozen at low passage numbers. We also used these cells as a base for the creation of a stable line expressing LifeAct-EGFP (see below). Both were cultured in Dulbeccos Modified Eagle Medium (DMEM) with GlutaMAX (#61965026, Thermofischer) supplemented with fetal bovine serum (FBS, 10% of final volume, #S1810-500, Biowest, France) and penicillin-streptomycin (1% of final volume, #15070063, Thermo Fisher, USA). Cells were maintained at 37°C with 5% CO2 in a humidified incubator. They were passed every 2 to 3 days with 1/4 to 1/10 dilution, using TrypLE (#12605036, Thermo Fisher, USA) so as never exceeding 80% confluence.

Both 3T3 fibroblast cell lines were amplified and frozen at low passage numbers. Cells were frozen in 1mL aliquots of roughly 2 millions cells/mL. To do so cells were detached using TrypLE (#12605036, Thermo Fisher, USA), centrifuged in culture medium and the pellet was suspended in a mix of fetal calf serum (50%), culture medium (45%) and DMSO (5%). They were conserved at -150°C and thawed when needed. The total number of passages was always kept below 20, and rarely went above 10 throughout this work. Cells were routinely tested for mycoplasma contamination.

##### ii. HeLa Fucci

Human cervical adenocarcinoma HeLa cells stably expressing FUCCI plasmid (gift from Buzz Baum lab) were cultured in DMEM GlutaMAX medium (Gibco, #61965-026) supplemented with 10% FBS (Gibco) and 1% penicillin-streptomycin (Gibco, #15140-122) at 37°C and 5% $CO_2$.

##### iii. MDCK

Madin Darby Canine Kidney (MDCK) type II were cultivated in DMEM low glucose media (#31885-023, Gibco) complemented with 10% FBS (#A5256701, Gibco) and 100 U/mL penicillin-streptomycin (#15140-122, Gibco). Cells were maintained at 37°C with 5% $CO_2$ in a humidified incubator. Cells were routinely tested for mycoplasma contamination.

##### iv. Dicty

The data on Dictyostelium discoideum are from the study[1], which was previously conducted in our laboratories. Dictyostelium discoideum, strain Ax3, were acquired from DictyBase.org and grown at 21°C in HL5 medium (#HLF3 Formedium). No new experiments were conducted for this study.

##### v. Primary DC

The data on primary DCs are from the study[1], which was previously conducted in our laboratories and specifies the collection conditions. No new experiments were conducted for this study.

#### 2. Viral transduction of LifeAct-EGFP

A stable cell line of 3T3 ATCC-2023 fibroblasts expressing LifeAct-EGFP was produced by lentiviral transduction followed by FACS sorting. Briefly, 4 µL of Lipofectamin 2000 (Thermofischer) were added in 200 µL of OptiMEM (Thermofischer) and supplemented with 2 µg of DNA, including:

- 0.3 µg of pMD2.G envelope protein (gift from Francois-Xavier Gobert, Institut Curie)
- 0.8 µg of PsPax2 reverse transcriptase, capside, integrase (gift from Francois-Xavier Gobert, Institut Curie)
- 0.9 µg of the plasmid of interest pLenti Lifeact-EGFP BlastR (Addgene Plasmid #84383)

1.6 million HEK-293FT cells (gift from the lab of Nicolas Manel) were plated in 2 mL of DMEM (#61965059, Thermofischer) and supplemented with the mix. 4 hours later the medium was replaced with 3 mL of the 3T3 culture medium. The next day viruses were harvested by collecting the medium and filtering it with a 0.45 µm membrane. 1 mL of virus solution was added to a well of a 6-wells plate containing cultures of 3T3 ATCC fibroblasts around 25. The third day 3T3 cells were washed twice with PBS and the medium was renewed. In the following days cells were amplified and sorted using FACS to collect only cells within a narrow window of LifeAct-EGFP expression.

#### 3. Magnetic Beads Preparation

Superparamegnetic Dynabeads M-450 Epoxy (#14011 Dynal, Thermo Fisher, USA) were chosen for the Magnetic Pincher technique as they ally several crucial features:

- Easy to localize and track in bright field using a defocusing microscopy approach
- Relatively easy to take up for cells (can be tuned by coating)
- Very monodisperse in diameter (see batch size measurement below)
- Able to acquire high magnetic moment magnitudes, thanks to a high density of magnetic elements[2] and a relatively large volume

The preparation of these beads for experiments comprised several steps. Bead size and magnetization measurements were conducted once per newly acquired batch of beads, in order to correct for batch-to-batch variability in these two quantities. Bead rinsing and coating were performed regularly on small number of beads, and coated beads were used in experiments within a few weeks. These procedures are detailed below.

##### i. Batch size measurement

For each batch of beads, we measured precisely the average diameter, using images of long chains of beads.

Briefly, a 35 mm glass-bottom petri dish was coated with 1% bovine serum albumin (Merck, Germany). A solution with the beads of interest in PBS (200'000 beads/mL) was added to the dish (≈ 2 mL). This dish was imaged (transmitted light, objective 100X NA = 1.4) under a uniform magnitude field of magnitude 5 mT. The beads self-organized to form long chains of more than 10 beads. (Methods Fig 1A). Approximatively 20 of these chains were imaged. Using the bead center detection routine detailed below, we determined to distance between all pairs of neighbors in the chains. The average was taken as the central tendency for the bead diameter in a given batch [See Methods Fig 1B and Table 1].

#### ii. Batch magnetization correction

The bead magnetization $M$ (A/m) as a function of the external field $B$ (T or mT) is known from the literature[2]. We fitted it with an empirical function of this form:

$$M(B) = k_{corrMag} \times K_{beads} \times \frac{aB^3 + bB^2 + cB}{\alpha B^2 + \beta B + \gamma} \qquad (Eq.1)$$

The values of parameters $a$, $b$, $c$, α, β, γ are fixed for all bead types. The value of $K_{beads}$ is adjusted for each bead type and depends of the density of magnetic particle in the beads. To account for batch-to-batch variability, the parameter $k_{corrMag}$ is adjusted for each new batch of beads purchased.

Briefly, a small chamber containing a very dilute suspension of beads, was placed in a gradient of magnetic field, using a single electromagnetic coil. The motion of these beads was filmed. In the regime of constant velocity, the force balance projected on the x-axis gives:

$$F_{drag} = F_{mag}$$

The magnetic force experienced by the bead due to its magnetic moment and the external gradient of field is balanced by its viscous drag (Methods Fig 1C). Being at low Reynolds number, this viscous drag is given by Stokes Law: $F_{drag} = 6\pi\mu R v_x$, where μ is the viscosity of water, $R$ the bead radius and $v_x$ the velocity of the bead along the x-axis. Once projected, the magnetic force is:

$$F_{mag} = m \cdot dB/dx = V \cdot M(B) \cdot dB/dx$$

Hence the relation:

$$M(B) = \frac{F_{drag}}{V \cdot dB/dx} = \frac{6\pi\mu R v_x}{V \cdot dB/dx} \qquad (\mathrm{Eq.}2)$$

With a gauss-meter (#GM08, Hirst Magnetics), the magnetic field and field gradient produced by the coil are measured, as a function of the supplied current intensity: the local gradient of magnetic field at the center of the setup is proportional to the field magnitude: $\frac{dB}{dx}|_{x=0} = k \cdot B(x=0)$. Beads in suspension in water were moved by the magnetic force, and this motion was filmed for several values of intensity. As shown on the kymograph on Methods Fig 1C, a constant velocity regime was reached, and $v_x$ was measured. Since $R$, $V$ and $dB/dx$ are known, an experimental $M(B)$ curve is obtained. By fitting it with [Eq. 1], the parameter $k_{corrMag}$ was adjusted to obtain the relation that will characterize the magnetization of one batch of bead (Methods Fig 1D).

| **Batch of beads** | **Coating** | **Diameter [nm] (mean ± std)** | **Coefficient $k_{corrMag}$** |
|---|---|---|---|
| M450-2023 | Fibronectin | 4477 ± 18 | 1.023 |
| M450-2025 | Fibronectin | 4493 ± 29 | 0.969 |
| M450-Strept | mPEG-biotin | 4506 ± 17 | 1.056 |

***Methods Table 1** – Calibration of superparamagnetic beads batches used in this study*

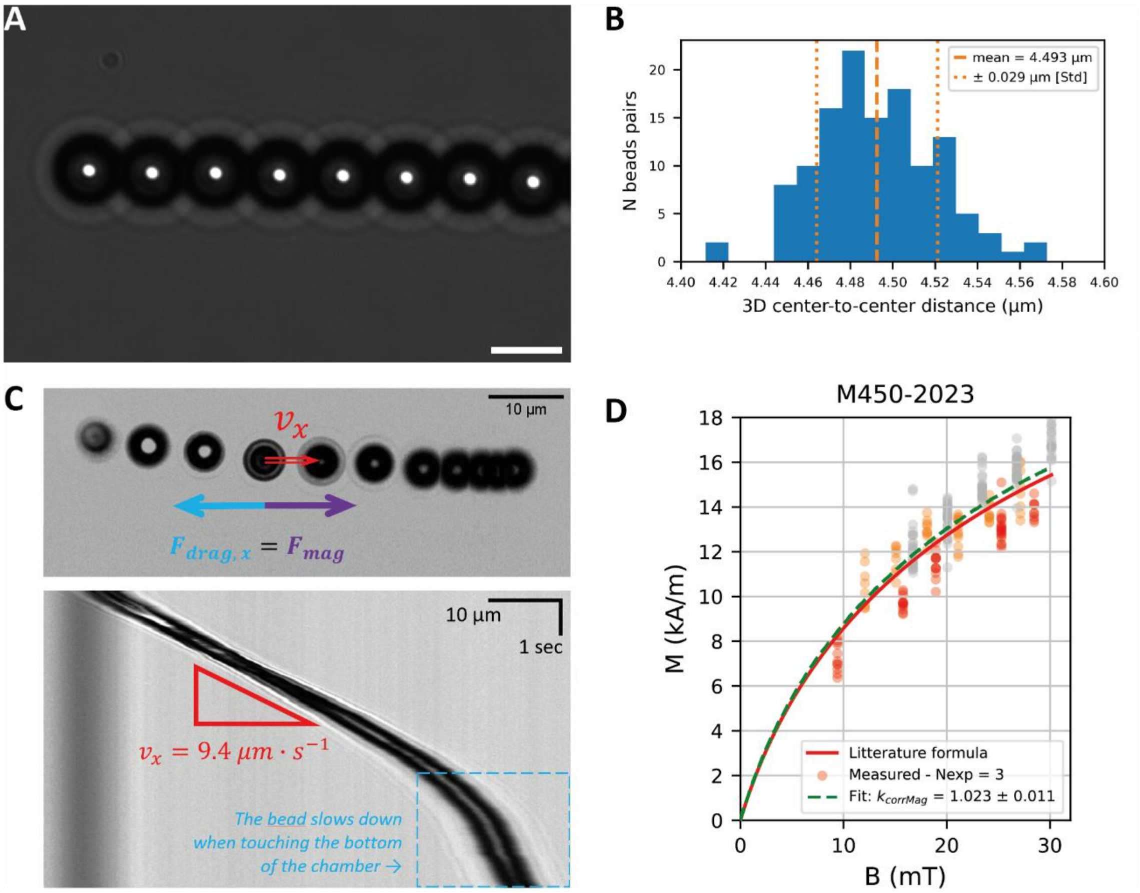


***Methods Figure 1 – M-450 Dynabeads properties measurement***

*A – A chain of M-450 Dynabeads in the configuration for the measurement of the average center-to-center distance. Scalebar 5 um. B – Distribution of center-to-center distances for one batch of M-450 Dynabeads (here the M450-2025, coated with fibronectin). N = 125. C – Typical images from a magnetization measurement experiment. On top: minimum projection of a film showing one M-450 Dynabeads moving in a magnetic field gradient while in free-fall. Bottom: the corresponding kymograph, along a line following the bead's trajectory. The speed is deduced from the angle of the dark line corresponding to the bead motion (excluding the part where the beads slow down as it contacts the surface). D – Typical magnetization calibration curve (here for the M450-2023, coated with fibronectin). Eq. 2 is used to compute M for each filmed bead for several values of B and dB/dx (which are proportional to one another in our protocol). The three colors of points represent each a replicate of the experiment, the red line corresponds to Eq. 1 with kCorrMag = 1 and the green line is the same Eq. 1 after the fit of kCorrMag.*

##### iii. Beads cleaning and coating

*Beads rinsing.* M-450 Dynabeads are conserved in distilled water with a concentration of $4 \cdot 10^8$ beads/mL. First, 30 µL of the stock solution were rinsed thrice with PBS using a magnet to retain the beads when removing the liquid phase. Then, two different coating protocol were applied.

*Coating with fibronectin.* A fibronectin solution is prepared by diluting 10 µg/mL fibronectin (#F1141, Sigma-Aldrich) in $NaHCO_3$ buffer pH 8.3 and filtering it with a 0.2 µm membrane. 100 µL of the solution is added in the aliquot containing the rinsed pellet of beads, and the mix is homogenized with a vortex for 30 seconds. The aliquot is placed on a rotating wheel for 3 hours so the beads get coated by the fibronectin.

*Coating with PEG*. Identical to fibronectin coating protocol, but replacing M-450 Epoxy by M-450 Dynabeads functionalized with streptavidin [included in the CELLection Biotin Binder Kit, #11533, Thermo Fisher, USA] and replacing the complete medium by 100 µL of mPEG(5K)-Biotin solution [#JK_A3097, Merck, Germany, 1 mg/mL in HEPES 10 mM, pH 7.4].

In both cases, the resulting concentration expected in the aliquot is 1 to $1.2 \cdot 10^6$ beads/uL. Aliquots were conserved at 4°C, with the cap wrapped in Parafilm, for up to one month.

#### 4. Micropatterning

We created arrays of fibronectin discs with 20 µm diameter on a PLL-PEG passivated glass coverslip. Fibronectin discs were disposed on a square lattice with a center-to-center distance of 90 µm. It was largely adapted from the method detailed in[3,4].

##### i. Photomask micropatterning

*Photomask design*. The masks geometry was designed with the software CleWin5. Quartz masks were purchased from JD-Photodata (UK).

*Coverslip passivation with PLL-Peg*. Round glass coverslips (diameter 25 mm, thickness #1) were cleaned with ethanol and exposed to plasma for 2 minutes. They were placed over 50 µL droplets of a 0.1 mg/mL PLL-Peg solution [PLL(20)-g[3.5]-PEG(2) from SuSoS, Switerland, diluted in HEPES 10 mM pH 7.4, and filtered with a 0.45 µm membrane]. They were incubated with the solution for 40 minutes, then removed, rinsed with milliQ water and gently dried with a tissue.

*Micropatterning with deep-UV*. The mask was cleaned with isopropanol, then the metallic side was exposed to deep-UV for 10 minutes. We used a UV lamp with wavelength λ = 254 nm and power P = 7 mW/m² (UVO Cleaner, Jelight). For each coated coverslip, a drop (~10 µL) of milliQ water was placed on the metallic surface of the mask. Coverslips were placed on top of these droplets, coated side down, without trapped air bubbles. The excess of water was removed by blowing compressed air from the top. The mask was exposed again to deep-UV-light for 10 minutes, this time with the non-metallic face toward the UV source. Finally, the coverslips were detached from the mask by pouring milliQ water, and dried.

##### ii. Chamber assembly

*Experimental chamber manufacturing*. A central 20 mm hole was cut in the plastic bottom of 35 mm petri dishes with a laser cutter (Epilog Laser, USA). Non-toxic silicon glue (Silicone SA 500, Zolux, France) was used to attach the patterned coverslip to the cut petri dish. The glue was left to dry for at least 4 hours and chambers were conserved for up to one week.

*Fibronectin addition.* This final step was performed on the day of an experiment, due to the lesser stability of fibronectin coating. Patterned dishes were incubated for 30 minutes with 125 µL of fibronectin solution [10 µg/mL fibronectin (#F1141, Sigma-Aldrich) in NaHCO3 buffer pH 8.3, filtered with a 0.2 µm membrane], using parafilm discs to spread the solution on the glass surface. The dishes were then rinsed twice with PBS. To visualize the patterns we occasionally added 4 µg/mL Alexa Fluor-conjugated fibrinogen (#F35200, ThermoFischer) to the solution.

#### 5. Magnetic Pincher Set-up

To monitor the actin cortex thickness in time, a pair of beads pinching the cortex must be tracked in 3D. This is done by acquiring a time-lapse movie of the beads illuminated in bright field, with the focus on the light spot below the beads.

In order to apply a controlled force on the cortex through the beads, a uniform magnetic field should be applied on the chamber. We devised a system to apply a programmed sequence of magnetic field, and synchronize the field generation with the image capture, so each image is associated with a timestamp and the exact magnetic field magnitude at this time.

##### i. Magnetic field generation

We used electromagnetic coils to generate a uniform magnetic field over the experimental chamber. A pair of such coils is positioned symmetrically along a common axis around the sample (pseudo-Helmoltz coils). When the same current circulates through both coils in the same direction, a quasi-uniform magnetic field in generated in the space between the coils. The magnitude of this field is proportional to the intensity of the electric current. In practice, the two coaxial coils (custom made by SBEA Technologies, France) are completed with a mu metal core (750 spires; length: 40 mm; inner diameter: 46 mm; outer diameter: 86 mm, see Methods Fig. 2) to increase the generated field. The coils are connected in series and powered by a bipolar operational power supply amplifier 6A/36V (Kepco, USA) controlled by the computer through a data acquisition module (National Instruments, USA). The maximum field generated is 55 mT (which correspond to the maximum supplied current, 6 A) with a gradient less than 0.1 mT/mm over the sample.

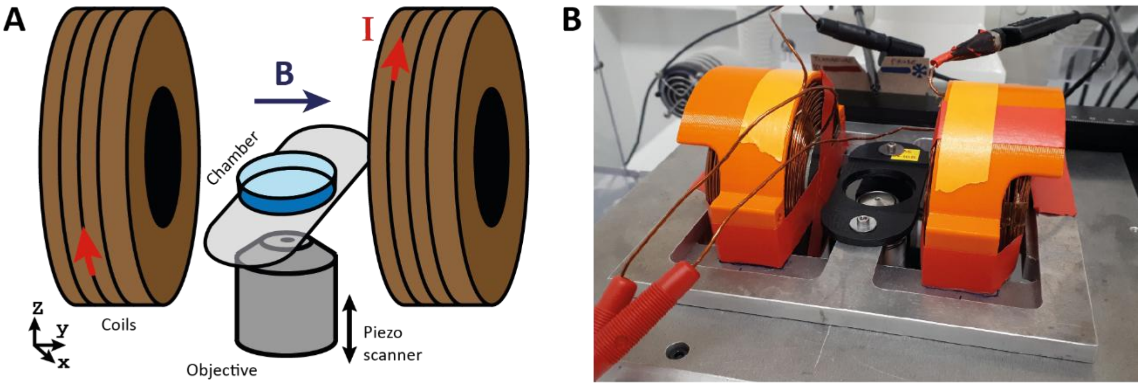


***Methods Figure 2 – Magnetic Pincher core set-up***
***A*** *– Core elements of the Magnetic Pincher set-up.* ***B*** *– Photograph of the actual configuration of the microscope stage, without the experimental chamber. The custom made structure supporting the dish holder enable the user to move the sample in the XY-plane while the coils position stays fixed.*

##### ii. Imaging system

In addition to the coils generating the magnetic field, the imaging setup includes the following elements:

1. Axio A1 inverted microscope (Carl Zeiss, Germany) with an oil-immersion 100× objective with NA = 1.4.
2. Orca Flash4 camera (Hammamatsu, Japan) with a 16-bits dynamic range.
3. PIFOC focus scanner to control the position of the focal plane along the Z-axis (PIFOC P-721.CDQ, Physik Instrumente, Germany).

4. Image acquisition software which can control both the camera and the Z-axis, and synchronously read the intensity sent to the coils at the time of capture of each image. We developed a Labview software, which we combined with a DAQ (data acquisition module, #NI6343, National Instruments, USA).
5. An environment control chamber to maintain the temperature of the sample to 37°C throughout the experiment (The Box & The Cube, Life Imaging System, Switzerland).

#### 6. Measure of cortical mechanics

Overall, the protocol consists in three phases: first, material preparation (a few days before the experiment): making the patterned chambers and incubating cells with beads to ingest. Second, on the day of the experiment, seed the cells in patterned chambers, let them adhere and add external beads (and the drug treatment if required). Third, bring the chamber to the microscope and run the custom Labview program to film beads pinching a cell's cortex while controlling the magnetic field.

The following sections detail the protocol in the case of 3T3 fibroblasts and HeLa FUCCI cells. The adjustments in the case of MDCK cells are presented at the end.

##### i. Cells preparation

*Incubation of cells with beads*. 48h hours before the experiment, approximately $2 \cdot 10^5$ cells are seeded in a 25 cm² culture flask with culture medium. 6 µL of fibronectin-coated M-450 Dynabeads solution are mixed in 1 mL of warm culture medium. This suspension is added to the culture, then the flask is gently rocked to homogenize the distribution of beads.

*Cells adhesion on micropatterns*. Cells are detached with TrypLE (#12605036, Thermo Fisher, USA) and resuspend them in warm imaging medium [Culture medium supplemented with 20 mM sterile HEPES buffer (#H_0887, Merck, Germany)]. The quantity of medium is adjusted to get approximately $1.5 \cdot 10^5$ cells/mL. 2 mL of cell suspension are transferred in a micropatterned chamber. The chamber is placed for 20 min in the incubator (37°C, 5% $CO_2$) so the cells start adhering on the patterns. Then, to remove the excess of non-adherent cells, the bottom of the chamber is washed with warm imaging medium, without ever drying the coverslip. The chamber is again incubated for 2 hours (37°C 5% $CO_2$) so that the cells adhesions mature to a steady state.

*Outer bead addition (without drug treatment)*. 2.5 µL of the coated beads solution were added in 0.5 mL of warm imaging medium. The mix was homogenized and added to the experimental chamber.

##### ii. Drug treatments

For all experiments involving chemical treatments, the drug was added with the outer beads as detailed below, instead of the normal procedure.

*Outer bead addition (with drug treatment).* The volume of medium in the experimental chamber to was adjusted to be 1 mL. The chosen drug was diluted at twice the target concentration in an aliquot containing 1 mL of warm imaging medium and 2.5 µL of the coated beads solution. This aliquot was vortexed and added to the experimental chamber 30 minutes before imaging. The references and concentrations of drugs used are detailed in Methods Table 1.

| Drug name | Concentration | Reference | Solvent used |
|---|---|---|---|
| Y-27632 (Y27) | 20 µmol/L | #1254 Tocris MilliQ | Water |
| LIMKi3 | 20 µmol/L | #435930 Sigma-Aldrich | DMSO |
| CK666 | 50 µmol/L | CK 666, #3950 Tocris | DMSO |
| Latrunculin A (Lat A) | 0.5 µmol/L | #L5163 Sigma-Aldrich | DMSO |
| DMSO | - | #D2438 Sigma-Aldrich | - |

***Methods Table 2*** *– References and concentrations of chemical drugs.*

##### iii. Depthograph acquisition

One or more beads were positioned within the field of view, ensuring that the beads were clearly separated from one another and remained stationary throughout the acquisition. The focal plane was initially positioned at the equatorial plane of the beads.

For each field of view, a Z-stack was acquired over a total axial range of 8 µm, with 401 frames and a 20 nm step size. Axial displacement was controlled using the piezoelectric actuator to ensure accurate and reproducible positioning along the Z-axis. This acquisition procedure was repeated until a minimum of eight Z-stacks of beads had been obtained.

##### iv. Measure of cortical mechanics

The electromagnetic coils power supply was switched on, and the magnetic field magnitude was set to a resting value (5 mT for 3T3 fibroblasts and HeLa cells, see below for MDCK cells). Then a cell whose cortex is pinched by a pair of beads is positioned in the field of view. We excluded beads in the following cases:

- The pair of beads is far from aligned with the magnetic field direction (angle > 30°).
- The light spot below the beads is hindered by extra or intra-cellular objects.
- Beads pinching the cortex have 3 nearest-neighbors or more. Such geometry makes the magnetic field in the region of the beads impossible to define properly.

We then acquired a time-lapse of these beads. It consists either in an observation under constant magnetic field, or in a series of compressions.

*Constant field experiment*. 5 to 10 minutes, one Z-stack of images every 600 ms, for a total of 200 to 1000 time-points. Each Z-stack consists in 3 slices, with a step of 0.5 µm, ideally centered on the plane of maximum intensity and with a delay of 50 ms between each step of the Z-stack.

*Compression experiment.* Series of 5 to 10 sequences comprising a resting phase, a compression phase where the field is brought to its maximum value, a relaxation phase where the field is brought back to its resting value, and another resting phase. See the Method Table 2 table] for precise phase durations and magnetic field magnitudes.

A given chamber was never imaged for longer than 2 hours. Typically, 10 to 20 cells could be acquired in each chamber, depending on the duration of the acquired time-lapse movies.

The setup produces two types of raw data. First, films of beads pinching cells, which are used to compute the beads position and eventually the cortex thickness over time. Second, data files containing for each image of each film: a precise timestamp, the value of the applied magnetic field, the position along the z-axis. To ensure a precise correspondence between the images in the film and the associated data in the table, the camera sends a trigger signal to the DAQ module every time an image is captured, and the module acquire in response the numeric data listed above.

| Phase | Resting Phase 1 | Initial relaxation | Constant Field | Indentation | Relaxation | Resting phase 2 |
|---|---|---|---|---|---|---|
| Duration (s) | 6 | 2 | 2 | 1.5 | 1.5 | 6 |
| B(t) | constant | sigmoid | constant | $\propto t^2$ | $\propto t^2$ | constant |
| B initial (mT) | 5 | 5 | 2 | 2 | 55 | 5 |
| B final (mT) | 5 | 2 | 2 | 55 | 5 | 5 |
| Z-stack | Yes, 3 slices, $\delta z$ = 0.5 µm | No | No | No | No | Yes, 3 slices, $\delta z$ = 0.5 µm |
| Delay between frames (ms) | 50 ms within a stack, 500 ms between stacks | 100 | 100 | 12 | 100 | 50 ms within a stack, 500 ms between stacks |

***Methods Table 2** – Details of the parameters used to control the setup during cortex indentation experiments*

#### v. Protocol adjustments for MDCK cells

*Cells preparation.* Cells are incubated overnight in a T25 flask with $\approx 6 \cdot 10^6$ M-450 Dynabeads coated with fibronectin. MDCK with internalized beads are detached with trypsin (#25300054 Gibco) for 10-15 min and then diluted in culture medium. To enrich the proportion of cells with internalized beads, suspended cells are disposed in a falcon vertically suspended above a magnet. After 20 s, the upper fraction of suspended cell is removed. About $2 \cdot 10^5$ cells from the remaining enriched fraction are added on coverslips with 20 µm diameter disk micropatterns. After 20 min incubation (37°C 5% $CO_2$), cells were attached to the substrate. Unattached cells were washed using DMEM-Fluorobrite (#A1896701 Gibco) supplemented with 20 mM HEPES (#15630-056 Gibco) and 100 U/ml penicillin-streptomycin, without drying the substrate. Cells are incubated at least 4h in incubator with DMEM-Fluorobrite to let them spread. Before imaging, $1 \cdot 10^6$ of extracellular M-450 Dynabeads coated with fetal bovine serum are added to the dish.

*Measure of cortical mechanics.* With MDCK cells, a stronger magnetic field had to be applied to keep the beads associated in pairs and to indent the cortex. In order to achieve this with our setup, a pair of identical permanent magnets (Supermagnete, Germany) was added to the coil so the magnetic field in the center of the setup when no current is supplied is 32 mT. Then, the coils could be used as normal with a controlled current intensity to increase or lower this value. In summary, the measure of cortical mechanics in MDCK was done like for 3T3 and HeLa cells, except that the resting field was 32 mT, the low field before the indentation was 3.5 mT, the maximum field at the end of the indentation phase was 70 mT and finally the field was brought back to its resting value of 32 mT.

#### 7. Measure of cortical thickness and actin quantity

##### i. Setup modifications

In order to correlate measurements from the Magnetic Pincher with a metric of F-actin quantity, we developed a way to successively measure cortical thickness (with our usual protocol) and image the fluorescence intensity at the cortex. To do so we adapted the Magnetic Pincher system on a spinning disc microscope (Cell Discoverer SD, Zeiss, Germany). The setup was functionally identical to the one described above.

##### ii. Experimental method

The goal of this protocol was to measure both the cortex thickness and fluorescence intensity at the cortex for a given time point. To do so, we needed to acquire Z-stacks of beads pinching the cortex of a cell in bright field and spinning disc fluorescence, with shortest possible time delay in between.

First, LifeAct-EGFP expressing fibroblasts were prepared as normally for a Magnetic Pincher experiment. A micropatterned chamber with cells adhered on 20 µm fibronectin discs was placed in the imaging setup. The camber was exposed to a constant, uniform field of 5 mT to trigger the beads magnetization.

Then, we acquired Z-stacks in both bright field and fluorescence channel of beads pinching a cell cortex. The stacks consisted in 9 frames with a step of δz = 400 nm. Images in both channels were acquired within 4 seconds, and such sequence was repeated every 30 seconds for 3 to 10 minutes. We also set a systematic offset of $\Delta Z_0$ = +4 µm on the fluorescence channel compared to the bright field channel. This way, when the latter is showing the light spot below beads, the former shows the contact region between the beads. Indeed, in a Magnetic Pincher experiment, we normally image planes which are below the beads, where the bright spot forms and in the case of this imaging setup, we measured a distance of ≈ 4 µm between the equatorial plane of a bead and the location of the brightest spot below it.

The results of this method were similar to those of a typical Magnetic Pincher experiment, except that the field was always kept constant at 5 mT, and that for each time-point, 9 z-positions were acquired (instead of 3 normally) for two channels (bright field and GFP).

### II – Image Analysis

Image analysis was performed using ImageJ[3] and custom scripts written in Python[5,6].

Code repository : https://github.com/jvermeil-biophys/CortExplore

#### 1. Bead Distance Computation

##### i. Tracking in XY

The in-plane position of each particle was determined using the spot of bright pixels around the bead center. By calculating the intensity-weighted centroid of its gray-level distribution (exactly as in Laplaud et al.[1]), the position of the centre of the bead can be determined with sub-pixel accuracy. More specifically, this approach resulted in an error of approximately 2 nm on the bead center localization, consistent with that obtained in previous in cellulo and in vitro studies[1,7,8].

##### ii. Tracking in Z with Depthographs

The localization of the beads along the Z-axis was done using the rich interference pattern that appears below a bead under incident light. As mentioned above, a "depthograph" – an average YZ-profile of the light pattern below a bead in bright field – was created for each experiment. Using several Z-stack of still, single beads (800 frames, step 20 nm), YZ-profile including the center of the bead in each frame are extracted. Then these profiles are aligned along the Z-axis using the point of maximal intensity as a reference, and averaged to obtain a depthograph (see Methods Figure 3, left part). A new depthograph was generated for every experimental chamber to account for small optical changes.

In order to compute the relative motion of beads along the z-axis in our films, an intensity profile along the y-axis through the bead center was compared with the depthograph (see Methods Figure 3, left part). By finding the row of the depthograph with minimal difference with that profile, we compute the position of the bead along z. The difference is calculated through a cost function which is the sum of the squared difference between pixel intensity. Using the depthograph as a common reference, we then compute the distance between any two beads along z. During the constant field part of the compression experiments, we acquired 3 images at 3 different z (500 nm apart) in a quick succession. In this case, we simultaneously compared the three profiles to the depthograph to locate the bead position with a greater precision.

The precision on this procedure was estimated by cross-correlating independent depthographs and computing the average error. We found an error of approximately 40 nm when considering an image triplet, and 75 nm with a single image.

This leads to a typical error of 5 nm and 7 nm on the 3D distance between the bead centers, with image triplets and singlets respectively. This evaluation of the error does not include the uncertainty on the beads diameter (roughly 25 nm, see Methods Table 1) which is not present when considering a single pair of beads. When accounting for it, one gets a typical error of 26 nm and 27 nm (for image triplets and singlets respectively).

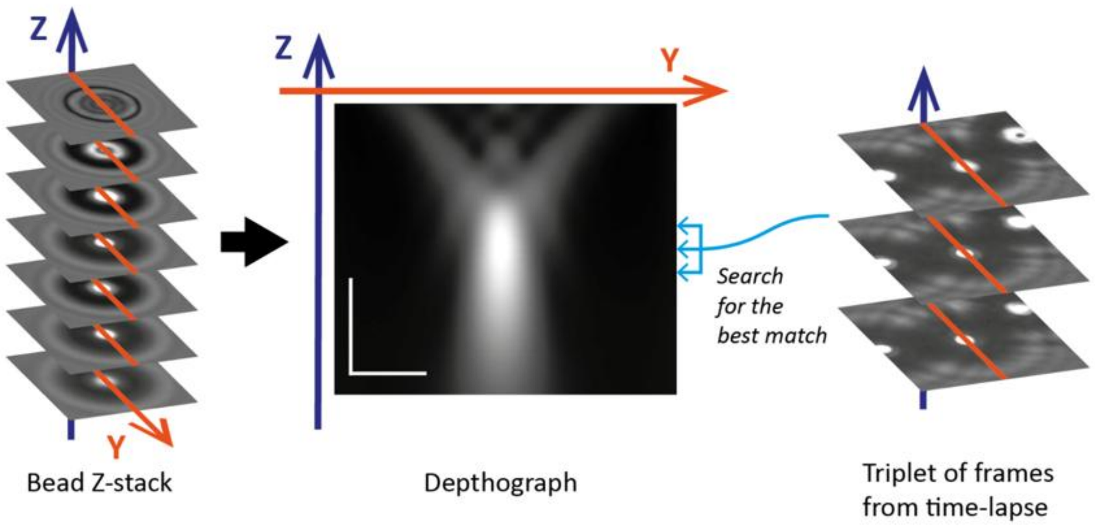


***Methods Figure 3 – Bead tracking along the Z-axis using a Depthograph.***

*On the left, illustration of a Z-stack used to generate the Dephtograph. We used stack of 400 images, vertically spaced by 20 nm. In the center, an example of such Depthograph, with the depth (Z) on the vertical axis and the profiles (orange lines) on the horizontal axis (vertical scale bar: 2 µm; horizontal scale bar: 1 µm). On the right, a typical application of the Depthograph: to locate the bead along the Z-axis, one can take its profiles (orange lines) on the frames of a Z-triplet, and compare them with every row of the Depthograph to find the best match. This approach, using the fixed distance between the 3 frames (0.5 µm) as an additional information, ensure the uniqueness of the best match and improve the precision. Therefore, each bead can be located within a common reference Depthograph and the distance Z between the beads pinching the cortex can be computed.*

#### 2. Magnetic Force Computation

##### i. Formula for a simple pair of beads

In order to compute cortex moduli, we need the expression of the pinching force, ie the force one bead on either side of the cortex applies on the bead on the other side. First, the total magnetic moment of a bead can be expressed as: $m(B) = V \cdot M(B)$ with $V = 4\pi(D/2)^2$ where $V$ and $D$ are the volume and diameter of the bead, and M(B) is the magnetization of the beads, measured experimentally.

The magnitude of the attractive force exerted by a bead on another is noted F and expressed as such:

$$F(B, d, \theta) = 3\mu_0 \cdot m_1(B) \cdot m_2(B) \times (3cos^2(\theta) - 1) \qquad (Eq. 3)$$

where $m_1(B)$ and $m_2(B)$ are the two beads magnetic moments, $d$ is the center-to-center distance, $\theta$ is the angle between the direction of the magnetic field and the center-to-center direction, and $\mu_0$ is the vacuum magnetic permeability (see Methods Fig. 4A-B). The factor $(3cos^2(\theta) - 1)$ expresses that this force is attractive of maximum magnitude when $\theta = 0$ (beads aligned with the field), and decreases as $\theta$ increases, to become repulsive when $\theta > 55°$. Note also that $F \propto d^{-4}$, meaning that the attraction decrease strongly as the distance increases.

Typical curves for $M(B)$ and $F(B)$ are shown on Methods Fig. 4C. The typical magnetic field range used in our experiments is 1 to 55 mT.

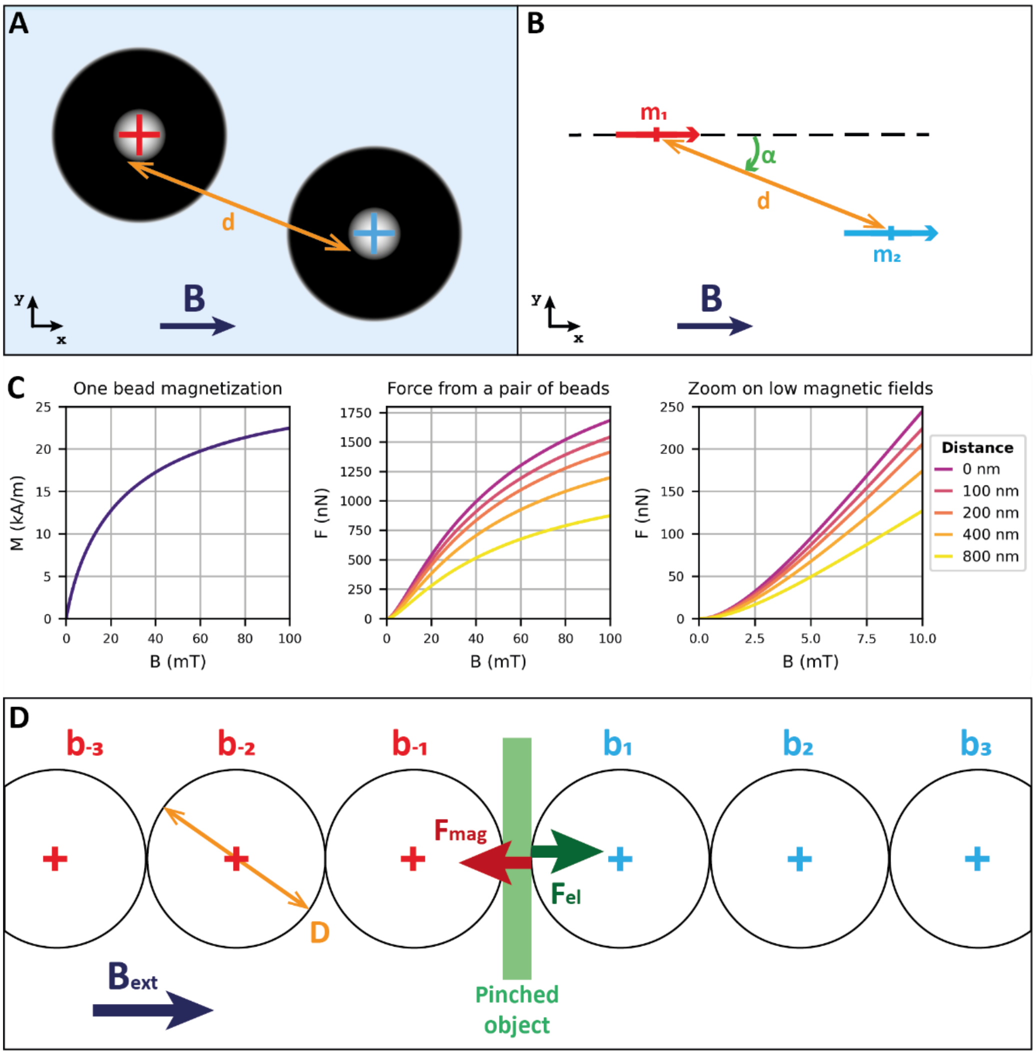


***Methods Figure 4 – Computing the force exerted by superparamagnetic beads***
***A, B*** *– Schematics illustrating the notations used in Eq. 3. In panel B the two beads are replaced by their equivalent punctual magnetic moment.* ***C*** *– Magnetization of M-450 magnetic bead and force exerted by a pair of such beads.* ***D*** *– Schematic illustrating the computation of added force in the case when the beads pinching an object have additional neighbors.*

#### ii. Corrections due to additional beads

When pinching the cortex with superparamagnetic beads, the situation was often more complex than a simple pair of beads because they tend to attract all their neighbors to form chains aligned with the magnetic field direction. Let's consider the case where the cortex is pinched by two arbitrary long chains of beads, represented on Methods Fig. 4D. The applied force is mostly due to the two beads in contact with the object, but the other beads of the chains also contribute, in two ways: increasing the local magnetic field, and attracting beads further away in the other chain. We assumed that a chain can be represented by series of punctual magnetic moments, aligned with the chain direction, and spaced apart by $D$ the beads diameter.

**Induced magnetic field.** In a chain, each bead is a magnetic dipole which generate its own magnetic field. It adds up with the external field to increase the magnetization of the neighboring beads. The expression of the field generated by a magnetic dipole along its axis is the following:

$$B(x) = \frac{\mu_0 \cdot m}{4\pi \cdot x^3}$$

Hence if a bead has one neighbor distant of one diameter $D$, the total field is:

$$B_{tot} = B_{ext} + B_{ind} = B_{ext} + \frac{\mu_0 \cdot m}{4\pi \cdot D^3} \approx B_{ext} + \frac{\mu_0 \cdot V \cdot M(B_{ext})}{4\pi \cdot D^3}$$

Here we assume that the neighbor magnetization is $M(B_{ext})$ and not $M(B_{ext} + B_{ind})$, hence neglecting a second-order contribution. Based on this formula, we chose to neglect the field induced by beads further than the first neighbors (at a distance 2D, the induced field is already 8 times lower).

In practice, when analyzing an experiment, we wanted to determine the field in the location of the two beads pinching the cortex. We considered only the two following cases:

i. The bead has only one neighbor, which is the other bead pinching the cortex. The correction applied was then:

$$\mathrm{Btot} = \mathrm{B_{ext}} + \mathrm{B_{ind}} = \mathrm{B_{ext}} + \frac{\mu_0 \cdot \mathrm{V} \cdot \mathrm{M(B_{ext})}}{4\pi \cdot (\mathrm{D} + \mathrm{h})^3}$$

ii. The bead has two neighbors: the other bead pinching the cortex, and a third bead on the other side. Then the correction was computed as:

$$B_{tot} = B_{ext} + B_{ind,1} + B_{ind,2} = B_{ext} + \frac{\mu_0 \cdot V \cdot M(B_{ext})}{4\pi \cdot (D + h)^3} + \frac{\mu_0 \cdot V \cdot M(B_{ext})}{4\pi \cdot D^3}$$

where $h$ is the measured cortex thickness.

**Added forces.** When an object is pinched between two chains, the total force applied is not only due to the two beads directly in contact, but also include a contribution from the other beads in the chains. With the notations of Methods Fig. 4D, the total magnetic force applied on the object results of the attraction of $(b_1)$ by $(b_{-1})$, plus the attraction of $(b_1)$ by $(b_{-2})$, by $(b_{-3})$, ..., plus the attraction of $(b_2)$ by $(b_{-1})$, $(b_{-2})$, $(b_{-3})$, etc. The attractive force between two beads that are very far apart is negligible, since it varies as $d^{-4}$. Thus, we chose again to disregard beads beyond the first neighbors.
In practice, when analyzing an experiment, we wanted to determine the total pinching force applied on the cortex. We considered only the three following cases:

i. There are only two beads pinching the cortex: $(b_{-1})$ and $(b_1)$. Then:

$$F_{tot} = F_{-1,1} = 3\mu_0 \cdot m_{-1} \cdot m_1 \times (3cos2(\theta) - 1)$$

ii. One of the two beads pinching the cortex has another neighbor: $(b_{-2})$ or $(b_2)$.

$$F_{tot} = F_{-1,1} + F_{-1,2} \ or \ F_{tot} = F_{-1,1} + F_{-2,1}$$

iii. Both beads pinching the cortex have another neighbor: we have $(b_{-2})$, $(b_{-1})$, $(b_1)$ and $(b_2)$.

$$F_{tot} = F_{-1,1} + F_{-1,2} + F_{-2,1}$$

with:

$$F_{-1,2} = \frac{3\mu_0 \cdot m_{-1} \cdot m_2}{4\pi \cdot (2D + h)^4}$$

and $F_{-2,1}$ has a similar expression, obtained by considering the relevant magnetic moments.

In order to simplify the calculations, the angle θ was taken to be zero in the expressions of the added forces ($F_{-2,1}$ and $F_{-1,2}$). Furthermore, only the magnetic moments of the two main beads $m_{-1}$ and $m_1$ were corrected with the induced magnetic field. Finally, beads further away in the chains were disregarded.

#### 3. Actin quantification

The analysis of these images is conducted in two parts:

1. Bright field images are analyzed with the usual 3D-tracking algorithm, which now performs the Z-detection of the beads positions based on 9 frames instead of 3; this results in a more robust detection of the beads position along the z-axis (described above).
2. Fluorescence images are analyzed with a custom-made python code, described below.

**Analysis of fluorescence images**. The raw experimental data consist in a TZYX-hyperstack — our successive z-stacks in the fluorescence channel. For each time point, the goal is to segment the cortex of the cell, determine which frame is the closest to the beads-cortex point of contact, and finally extract intensity profiles (perpendicular to the cell edge) to quantify the fluorescence intensity at the cortex. First, we detect the approximate location of the cell in each frame, by computing binary mask of the cell (thresholding, method isodata), and finding the largest circle which fit inside the mask (function `polylabel`, from the python module `shapely`). The we use this rough estimate of the cell shape (see Methods Fig. 5A-B) as the initial parameter for a custom-made cortex segmentation function using the Viterbi algorithm to control the contour regularization (see GitHub, see Methods Fig. 5C-E).

This way, actin intensity at the cortex location can be measured around the cell perimeter, as it is shown on Methods Fig. 5E. The fluorescence intensity over the cell surface can also be represented as on a planisphere (Methods Fig. 5F).

Importantly, this fluorescence intensity value has been normalized. This is crucial to be able to compare measures on different cells, as the level of expression in LifeAct-EGFP might vary. To do so we computed the average fluorescence intensity in the whole cell, minus the beads and the nucleus: $I_{cell}$. We also measured the background fluorescence far from the cell: $I_{bg}$. The normalized intensity at the cortex $I$ was computed as such from the raw value $I_{raw}$:

$$I = \frac{I_{raw} - I_{bg}}{I_{cell} - I_{bg}}$$

The next step is to determine the location of the beads-cortex contact point. Using the information we got from the 3D-tracking of the beads in the bright field channel, we compute the location of the beads-cortex contact point in the fluorescence stack. We finally extract profiles of fluorescence along lines that are perpendicular to the cortex around the contact point. Such profile is represented on Methods Fig. 5F. With this (normalized) intensity profile, F-actin quantity in the cortex is computed by fitting a Gaussian curve on a narrow zone around the peak of intensity. Said curve has the following expression:

$$G(x) = \frac{Q}{\sqrt{2\pi\sigma}} \exp\left(-\frac{(x-\mu)^2}{2\sigma^2}\right)$$

where the fitted parameters are μ the mean, σ the standard deviation and $Q$ the amplitude of the Gaussian. This parameter $Q$ constitutes our metric for F-actin quantity in the cortex.

In practice, we compute $Q$ in a small zone around the point of contact: we consider five angles (contact angle, contact angle $\pm$ 1°, contact angle $\pm$ 2°) in 3 planes ($Z_{contact}$, $Z_{contact}$ − 400 nm, $Z_{contact}$ − 800 nm) for a total of 15 measurements of $Q$. We remove outliers in this series of values and take the average to obtain the final result. Methods Fig. 5G shows an example of measures for one cell at 7 time points, as a function of the corresponding cortex thickness $H_{5mT}$ (computed from the matching bright field images).

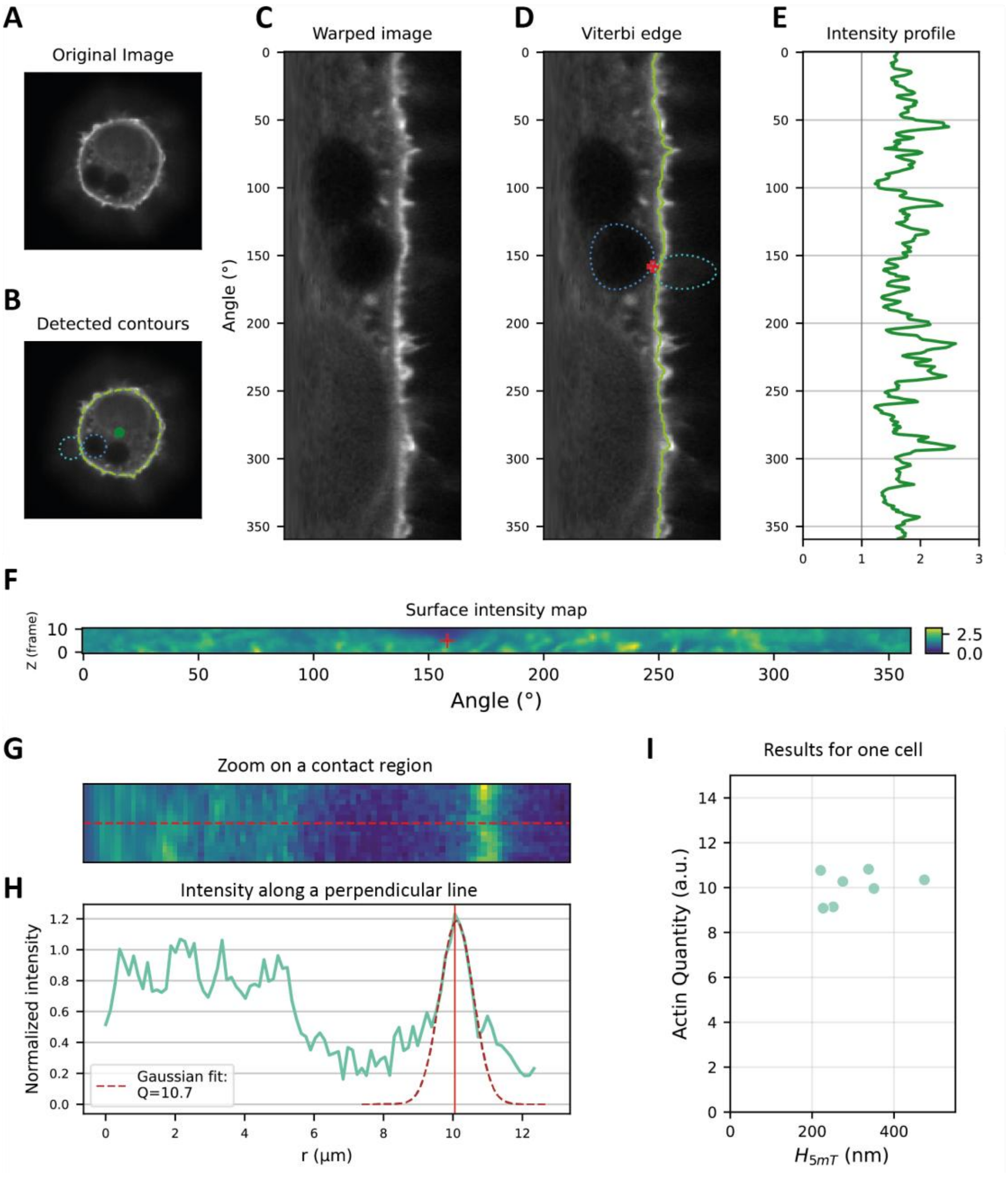

***Methods Figure 5 – Image analysis method of actin fluorescence quantification**. A – 3T3 fibroblast with F-actin labeled with LifeAct-EGFP, imaged with spinning disc confocal microscopy (100X). B – Same image, annotated with the approximate center of the cell (green dot), the computed contours of the beads (blue and cyan) and the detected contour of the cell (lime green). C – Warped version of image (A). D – Annotated version of (C), where are shown the (warped) contour of the beads (blue and cyan) and the detected contour of the cell (lime green). The red cross shows the point of contact as computed from the 3D tracking of the beads. E – Normalized intensity profile along the cell contour. F – Cell surface intensity map, where each row corresponds to the intensity along the cell contour in one z-plane. The red cross shows the point of contact. G – Zoom over a contact region. The red line is the radius going through the contact point. H. Intensity profile along the red line from (G). The vertical red line shows the maximum intensity at the cortex. The fitted brown curve is a Gaussian curve with an adjustable amplitude Q, which fits the profile over a narrow region around the maximum. I – Quantity of actin measured versus cortex thickness for one cell, imaged for 7 time-points. The cell is the same than for panels (G) & (H).*

### III - Models, Fits, statistics

Data analysis was performed using custom scripts written in Python[6,9–13].

#### 1. Physical model

##### i. Chadwick model

We consider the cortex as a thin layer, linearly elastic and isotropic, indented symmetrically by two large spheres with no contact adhesion. To account for this geometry, a model has been derived in 2002 by Chadwick[14], and revisited more recently [15]. It results in a force-indentation relation expressed as such:

$$F = \frac{\pi E R (H_0 - h)^2}{t \cdot H_0}$$

with $F$ the force applied by the beads, $h$ the thickness of the cortex (the two measured quantities), $E$ the cortex elastic modulus, $H_0$ its undeformed thickness (the two parameters to be fitted), and $R$ the radius of the beads. This can be read as a force-indentation relation since $\delta = H_0 - h$ is the indentation: here $F \propto \delta^2$.

The constant t depends on the material's compressibility: t = 3 for an incompressible material[14] and t = 2 if Poisson's ratio is zero (a more complete formula is found in[15]). More generally, any assumption regarding the compressibility of the material will result here in only a small change in a scaling factor. We used t = 3 throughout this work.

This result can be understood as a variation of the usual Hertz contact formula, where $F \propto \delta^{3/2}$. In this case, a sphere indents a semi-infinite medium and the deformed region grows both radially and in the depth of the material. Here, the layer is very thin compared to the spheres radius, and the deformed region grows only radially as the indentation increase: this geometric difference explains the new exponent in the scaling.

##### ii. Dimitriadis model

As an alternative to Chadwick's $F - \delta$ relation, we used another model in Supp. Fig. 11. This formula, proposed by Dimitriadis et al. in [16], is an expansion of the Hertz contact formula in the case of a sample of finite thickness indented by a sphere. The model assumes that the thickness of the indented layer $h$ is large compared to the radius of the contact surface $a = \sqrt{R\delta}$, with $\delta = H_0 - h$. As such, the formula introduces the quantity $\chi$ defined as:

$$\chi = \frac{a}{h} = \frac{\sqrt{R\delta}}{h} \ll 1$$

The complete formula is the following:

$$F = \frac{4E}{3(1-\nu^2)} R^{1/2} \delta^{3/2} \cdot [1 - A\chi + B\chi^2 - C\chi^3 + D\chi^4],$$

with the constants defined as:

$$A = \frac{2\alpha_0}{\pi},\ B = \frac{4\alpha_0^2}{\pi^2},\ C = \frac{8}{\pi^3}\left(\alpha_0^3 + \frac{4\pi^2}{15}\beta_0\right),\ D = \frac{16\alpha_0}{\pi^4}\left(\alpha_0^3 + \frac{3\pi^2}{5}\beta_0\right)$$

and:

$$\alpha_0 = -1.041,\ \beta_0 = 0.028$$

Interestingly, in their study, Dimitriadis and his co-authors notice that this formula gives surprisingly good results at high values of $\chi$ and even when $\chi \approx 1$. Therefore, when applying the model to our experimental $F - h$ curves we restricted ourselves to the parts of the curves where $\chi < 0.75$, and took off curves where this condition resulted in keeping less than 15 points. Doing so, we removed more than half the number of curves (414 over 1265, see Supp. Fig. 11), which is why we did not consider the Dimitriadis formula as a viable alternative to Chadwick's.

#### 2. Curve fitting

##### i. Fitting Chadwick F-$\delta$

**Fitting strategy.** The Chadwick $F - \delta$ relation is, in the case of our system:

$$F = \frac{\pi E R (H_0 - h)^2}{3H_0}$$

with $F$ the force applied by the beads, $H$ the thickness of the cortex (the two measured quantities), $E$ the cortex elastic modulus, $H_0$ its undeformed thickness (the two parameters to be fitted), and $R$ the radius of the beads.

However, in order to obtain $E$ and $H_0$ from our experimental data, it is more accurate to inverse the relation and treat $h$ as the Y-variable and $F$ as the X-variable. This is because $F$ is almost fully set by the applied external magnetic field (the center-to-center distance variations stay low, as the cortex is thin compared to the beads radii), and thus less affected by noise. Meanwhile, $h$ is a measured quantity and prone to measurement noise. As such, a least-squares fit is more accurate on a $h = f(F)$ relation rather than $F = f(h)$. Consequently, we used the inverted formula to fit cortical thickness $h$ as a function of applied force $F$:

$$h = H_0 - \sqrt{\frac{3H_0 F}{\pi E R}}$$

In order to fit $E$ and $H_0$ with this formula on the experimental $F - H$ indentation curves, we used the Python function `scipy.optimize.curve_fit,` with the following initial values and boundaries for the fitted parameters:

- Initial $H_0$: $H_{0,init} = max(h)$
- Initial $E$: $E_{init} = 3\, max(h) \cdot max(F) \;/\; \left[\pi R\left(max(h) - min(h)\right)^2\right]$
- Bounds for $H_0$ and $E$: $[0, +\infty[$

The outputs of this Python function were the values for the fitted parameters and the associated variance. We used the following formula to compute the 95% confidence interval for each fitted parameter:

$$Cihw \;=\; t(97.5, N-2)\ \backslash cdot\ \backslash sqrt\{Var\}$$

with $Cihw$ is the confidence interval half-width, $N$ is the number of experimental points and $t(97.5, N-2)$ is Student's t coefficient (for 97.5% on each side and N-2 degrees of freedom) and $Var$ is the variance of the fitted parameter.

**Curve validation criteria.** To be considered valid, a set of fitted parameters had to meet two criteria:

- On the coefficient of determination: $R^2 \geq 0.6$.
- On the Chi-squared coefficient: $\chi^2 \leq 1$.

The Chi-squared is defined as follow:

$$\chi^2 = \frac{1}{N-2}\sum\left(\frac{Y_{meas} - Y_{fit}}{errY}\right)^2$$

where $errY$ is the estimated error on the measured quantity $Y$. Here the $Y$ quantity is the cortical thickness $h$ and we assumed $errY$ = 7 nm, following the estimation of the errors detailed above.

##### ii. Fitting E-h relations

A simple linear regression using the ordinary least squares method on $E$ and $H_0$ gives unsatisfying results. Indeed, this approach implies that one of the two quantity is measured while the other is controlled and thus exempt of measurement errors: this is why for a set of paired values (X, Y), ordinary least squares will give different results when fitting X vs. Y and Y vs. X.

In our case both $E$ and $H_0$ are measured and affected by errors, thus we decided to use Deming regression, an errors-in-variables model. Deming regression assumes that the ratio of the errors' variance is known: $\delta = \sigma_y^2/\sigma_x^2$. In our case, while the measurement error on H could be computed, the error on E is difficult to estimate properly. Thus, we chose to define δ as the ratio of the variances of $E$ and $H_0$ themselves: $\delta = Var(E)/Var(H_0)$.

In practice, we used Deming regression implemented in Python in the library `odrpack`[13]

#### 3. Statistics

##### i. Significance Tests

To test for the significance of correlation between two variables, for instance for all representations of the thickness-modulus coupling, we used the p-value associated to the Pearson correlation coefficient. To compute it in Python we used `scipy.stats.pearsonr` with `alternative = 'two-sided'`.

To test if two non-normal distributions are significantly different from each other, we used the non-parametric Mann-Whitney-Wilcoxon test, implemented in Python as `scipy.stats.mannwhitneyu`.

To test if a set of measures was normally distributed, we used the Shapiro-Wilk test (in this test, the null-hypothesis is that the distribution is normal). To test if a set of measures was log-normally distributed we used the Shapiro-Wilk test after taking the logarithm of these measures. We used the Python implementation `scipy.stats.shapiro`.

##### ii. Weighted averages

For all computations of average elastic modulus per cell (for instance when showing the thickness-modulus coupling with one point per measured cell, inset of Fig. 2A) we used a weighted average. In this calculation, the weight of each value of $E$ was set to $E/Cihw(E)^2$, where $Cihw(E)$ is the confidence inverval half-width for the fitted value of $E$. This method was adopted because values of E are prone to be affected by noisy experimental $F - H$ curves. Thus, using weights that will sharply decrease as the confidence interval width for the fitted value of $E$ increases allow to softly filter the values of $E$ fitted from these noisy curves.

#### IV – Bibliography

10. Seabold, S. & Perktold, J. statsmodels: Econometric and statistical modeling with python. in *9th Python in Science Conference* (2010).

11. Waskom, M. L. seaborn: statistical data visualization. *Journal of Open Source Software* **6**, 3021 (2021).

12. team, T. pandas development. pandas-dev/pandas: Pandas. Zenodo https://doi.org/10.5281/zenodo.3509134 (2020).

13. Vale, H. odrpack.

14. Chadwick, R. S. Axisymmetric Indentation of a Thin Incompressible Elastic Layer. *SIAM J. Appl. Math.* **62**, 1520–1530 (2002).

15. Wu, J. & Ru, C. Q. Spherical indentation of an elastic layer on a rigid substrate revisited. *Thin Solid Films* **669**, 500–508 (2019).

16. Dimitriadis, E. K., Horkay, F., Maresca, J., Kachar, B. & Chadwick, R. S. Determination of Elastic Moduli of Thin Layers of Soft Material Using the Atomic Force Microscope. *Biophysical Journal* **82**, 2798–2810 (2002).
